# Essential functions of Inositol hexakisphosphate (IP6) in Murine Leukemia Virus replication

**DOI:** 10.1101/2024.02.27.581940

**Authors:** Banhi Biswas, Kin Kui Lai, Harrison Bracey, Siddhartha A.K. Datta, Demetria Harvin, Gregory A. Sowd, Christopher Aiken, Alan Rein

**Affiliations:** HIV Dynamics and Replication Program, National Cancer Institute-Frederick, P.O. Box B, Frederick, MD 21702-1201, USA; Department of Pathology, Microbiology, and Immunology, Vanderbilt University Medical Center, Nashville, TN 37232-3263, USA

## Abstract

We have investigated the function of inositol hexakisphosphate (IP6) and inositol pentakisphosphate (IP5) in the replication of murine leukemia virus (MLV). While IP6 is known to be critical for the life cycle of HIV-1, its significance in MLV remains unexplored. We find that IP6 is indeed important for MLV replication. It significantly enhances endogenous reverse transcription (ERT) in MLV. Additionally, a pelleting-based assay reveals that IP6 can stabilize MLV cores, thereby facilitating ERT. We find that IP5 and IP6 are packaged in MLV particles. However, unlike HIV-1, MLV depends upon the presence of IP6 and IP5 in target cells for successful infection. This IP6/5 requirement for infection is reflected in impaired reverse transcription observed in IP6/5-deficient cell lines. In summary, our findings demonstrate the importance of capsid stabilization by IP6/5 in the replication of diverse retroviruses; we suggest possible reasons for the differences from HIV-1 that we observed in MLV.

## INTRODUCTION

The orthoretroviruses are divided into six genera (alpha-, beta-, gamma-, delta-, epsilon-, and lenti-retroviruses). While the replication of viruses in different genera is similar in broad outline, there are many significant differences in the details. In the present work, we describe a feature of gammaretrovirus replication that differs from that of the lentivirus HIV-1, the best-studied retrovirus.

Retroviruses are initially assembled and released from virus-producing cells in the form of an “immature” particle. Immature particles are formed from ∼1500-2000 copies of the Gag polyprotein, and contain the viral RNA and other proteins, all enclosed in a lipid bilayer derived from the plasma membrane of the cell. After the particle has been released, it undergoes maturation, in which Gag is cleaved by the viral protease into a discrete series of cleavage products. These products always include the “capsid” protein (CA), which assembles within the free virion into a structure termed the “mature core” or “mature capsid”. This structure encloses the viral RNA along with reverse transcriptase (RT), integrase, and another Gag cleavage product, the nucleocapsid (NC) protein. In HIV-1, the mature core frequently assumes a conical shape ^1^.

In recent years, it has been recognized that a small molecule, inositol hexakisphosphate (IP6), contributes to the assembly of both the immature particle and the mature core in HIV-1 ^2–5^. IP6 is abundant in mammalian cell cytoplasm ^6^ and is packaged in immature HIV-1; its presence within the virion makes it available during assembly of the mature core. In the immature particle, Gag is principally arranged as a lattice of hexamers, and IP6 is coordinated within these hexamers by two rings of lysine side-chains in the capsid domain of Gag (i.e., the region of Gag that will give rise to CA upon maturation) in these hexamers ^4,7–10^. Following maturation, IP6 is localized near the N-terminus of CA in the mature core, now in association with basic amino acids in CA pentamers and hexamers. This association profoundly increases the stability of the core, and the optimum stability appears to be essential for the successful reverse transcription of the viral RNA into DNA as required for viral replication ^4,5,11^.

In the present work, we have investigated the role of IP6 in the replication of the gammaretrovirus Moloney MLV. We now report that IP6 is packaged in MLV, and in many respects, it contributes to MLV replication in close analogy to its roles in HIV-1. However, one notable difference is that the efficiency of MLV infection is reduced in target cells that are deficient in IP6 or related molecules; in contrast, infection by HIV-1 is unimpeded in these cells. We propose that this difference arises from a divergence between gammaretrovirus and lentivirus reproduction: gammaretrovirus cores must partially disassemble in the cytoplasm of the infected cell ^12–15^, while lentivirus cores evidently remain intact until penetrating into the nucleus ^16–18^. It seems likely that the encapsidated IP6 is lost upon the cytoplasmic dissociation of MLV cores, while it is retained in HIV-1 cores.

## RESULTS

### IP6 enhances endogenous reverse transcription in MLV

Reverse transcription during HIV-1 infection has been analyzed in great detail. Recent studies suggest that the viral DNA is synthesized within the mature core, which remains largely or entirely intact until it reaches the nucleus of the newly infected cell ^16–18^. Successful infection appears to require optimal stability of the core; the structure is maintained by interactions between its constituent capsid (CA) molecules, as well as by small molecules associated with it ^5,11,19^. IP6 has a significant impact on reverse transcription in these experiments: cores isolated from virions can synthesize viral DNA *in vitro* if they are provided with optimal concentrations of IP6. In turn, the IP6 requirement is related to the stability of the core, as the requisite IP6 concentration is inversely correlated with core stability in HIV-1 mutants ^11,20^.

We tested the ability of cores isolated from MLV particles to synthesize viral DNA (“endogenous reverse transcription” or “ERT”). We prepared virions composed of WT MLV proteins and packaging either WT MLV genomes or the genome of an MLV-derived vector encoding firefly luciferase (pBabeLuc). The particles were produced in transiently transfected 293T cells and partially purified by pelleting through a sucrose cushion. To measure their ERT activity, we permeabilized the virions with melittin ^11^, added dNTPs and other possible cofactors, and after incubation at 37°C for 10 hours, assayed the mixtures for luciferase DNA by qPCR. As shown in Figure 1A, we found that the ERT reaction was promoted by the inclusion of IP6 and rNTPs, in addition to the dNTPs used in DNA synthesis. The activity was also strongly stimulated by melittin and profoundly inhibited by AZTTP, as expected for a reaction catalyzed by MLV RT ^21,22^.

**Figure 1:**
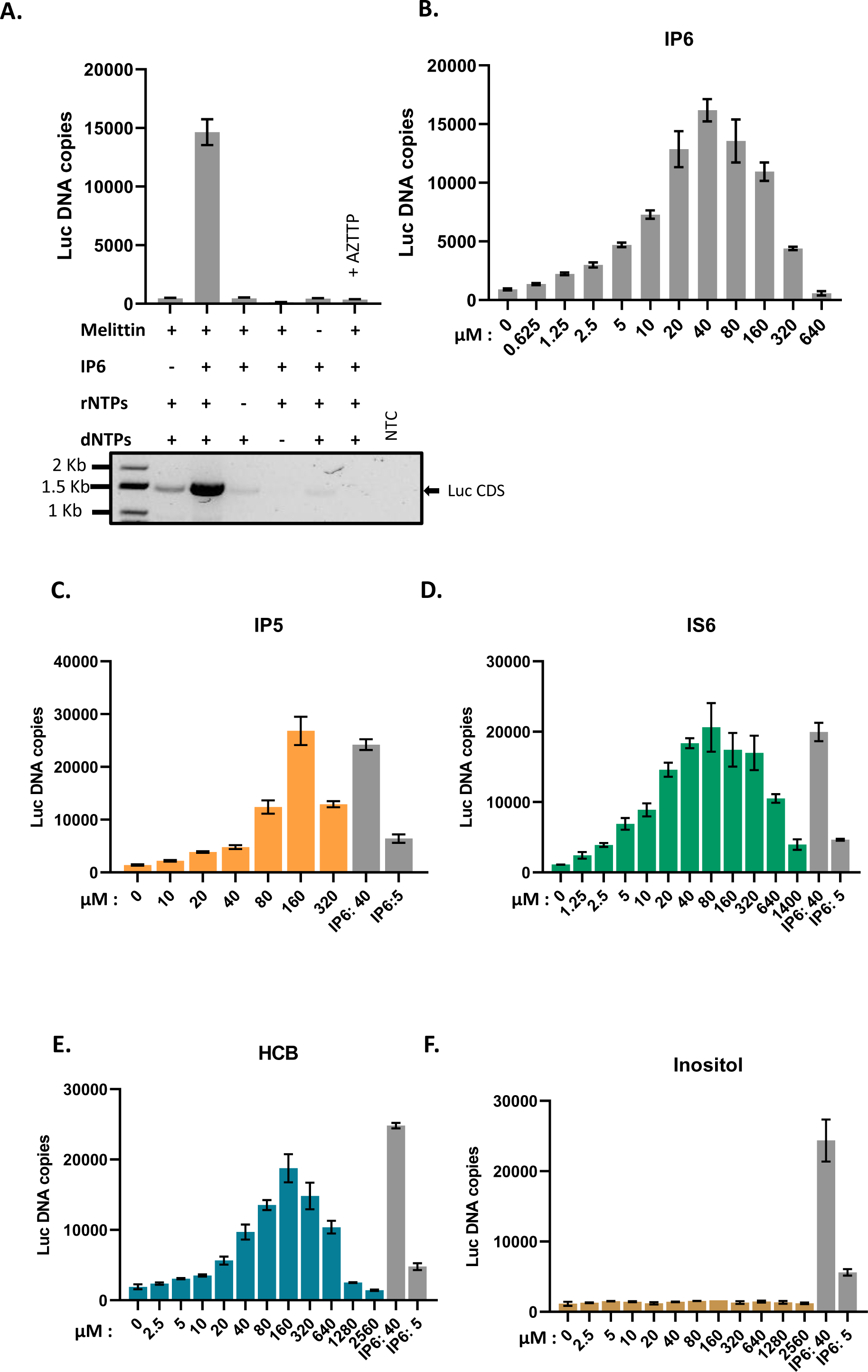
IP6 promotes ERT in MLV: A) The bar graph shows ERT product formation in the presence and absence of different reagents as indicated in the x-axis of the graph. The y-axis represents absolute copies of luciferase DNA reverse transcribed from the MLV-derived luciferase vector RNA packaged in MLV. In one sample, 30 μM AZTTP was included in the reaction mixture. Below the bar graph is an agarose gel showing amplification of a longer product (luciferase coding sequence, 1500bp) after ERT. NTC-no template control for the PCR reaction. B) IP6 titration in ERT assay. The graphs represent the mean ± SD of three replicates in the qPCR measurement of a single experiment selected from at least two independent experiments with similar results. C)-F): ERT product accumulation by titrating different potential co-factors in place of IP6: C) IP5, D) IS6, E) HCB (Mellitic acid), and F) Inositol.

As a further test of the ERT reaction, we performed PCR on the ERT products with a primer pair spanning the entire luciferase coding sequence (∼1.5 kbp). As seen in the lower panel of Figure 1A, the synthesis of the full-length luciferase DNA was almost completely dependent upon the addition of both IP6 and rNTPs, mirroring the qPCR results; it also required the melittin treatment and was sensitive to inhibition by AZTTP.

The reaction mixtures used above contained 40μM IP6 (where indicated), a concentration similar to that in mammalian cell cytoplasms ^6^. It was of interest to determine the ERT activities over a range of IP6 concentrations. As shown in Figure 1B, 40μM is indeed the optimal concentration under our ERT assay conditions, with significantly lower yields obtained at IP6 concentrations either ≤ 10μM or > 160μM.

The ERT documented above was dependent upon the inclusion of melittin in the reaction (Figure 1A). Titration showed that melittin was most effective at ∼3.13-12.5 μg/ml (Figure S1A), and 6.25 μg/ml was used in all subsequent experiments. We also found that melittin could be replaced by Triton X-100, with a threshold effective concentration of ∼0.2 mM (Figure S1B). The ability of this mild non-ionic detergent to replace melittin in ERT supports the hypothesis that melittin permeabilizes the viral membrane, presumably giving the RT and template RNA inside the core access to the dNTPs in the ERT reaction buffer, as required for DNA synthesis.

We also monitored the time-course of the ERT reactions in the presence and absence of IP6. These experiments used primers specific for either (-) strand strong stop DNA (“early”); DNA made following the first strand transfer (“intermediate”); or DNA made following the second strand transfer (“late”). Note that in the assays described above, only the luciferase sequences (made by reverse transcription of the pBabe-Luc luciferase vector in the virus preparations) were measured; in contrast, in this kinetic experiment, the primers amplified non-coding sequences present in both the luciferase vector and the intact MLV “helper” virus present in our virus preparations. As shown in Figure S1C, we found that all three regions of the viral genomes were produced, as expected; the stimulation by IP6 was particularly significant for the later ERT products. We also noted that the amounts of “early” and “intermediate” DNA decreased after the initial peak; it is conceivable that the DNase used in virus purification is responsible for this decline, but we have not investigated this point.

Further characterization of the ERT reaction is shown in Figure S1D. Interestingly, a 5-fold reduction in the concentration of the four rNTPs caused a profound decrease in ERT. Moreover, the specific omission of rATP, even in the presence of all the other cofactors, drastically impaired ERT product formation. Conversely, rATP alone could support ERT, even in the absence of other rNTPs.

In our standard ERT reaction buffer, rATP is at 6.7mM, whereas the other rNTPs are at significantly lower concentrations. To test the possibility that the apparent requirement for rATP is simply due to its uniquely high concentration in these experiments, we also tested ERT with the other rNTPs supplied singly at 6.7 mM. As shown in Figure S1D, rCTP, like rATP, supports full ERT activity at this concentration, whereas rGTP and rUTP do not. At present, we do not fully understand the rNTP requirement in ERT of MLV. It seemed possible that rNTPs fulfill the same function as IP6. As shown in Figure S1E, some ERT activity could be detected in the absence of rNTPs when the IP6 concentration was raised approximately 10-fold beyond the optimum in the standard reaction, but even this activity was far below that seen when rNTPs were included. The results suggest that rNTPs and IP6 are both necessary for maximal ERT activity.

One possible explanation for the apparent dependence on IP6 for ERT activity (Figure 1) could be that MLV RT requires IP6 for catalytic activity. We tested this hypothesis by lysing MLV particles with Triton X-100 and assaying, by product-enhanced reverse transcriptase (PERT), the ability of the released RT to copy an external template, i.e., MS2 RNA, into DNA by qPCR. As shown in Figure S1F, varying the IP6 concentration between 0 and 200 μM had no significant effect on the reaction. Thus, IP6 promotes ERT (Fig. 1A) by an effect on the cores, not on the RT enzyme *per se*.

### Other cyclic polyanions also promote ERT in MLV

We also tested other small molecules for their ability to replace IP6 in the ERT assay. As shown in Figure 2, we found that several cyclic polyanions, i.e., inositol pentakisphosphate (IP5) (Figure 1C), inositol hexasulfate (IS6) (Figure 1D), and hexacarboxybenzene (HCB or mellitic acid) (Figure 1E), could all promote ERT. Interestingly, IS6 was nearly as active as IP6, while IP5 and HCB were somewhat less active. In contrast, the uncharged cyclic molecule inositol was completely inactive (Figure 1F).

**Figure 2:**
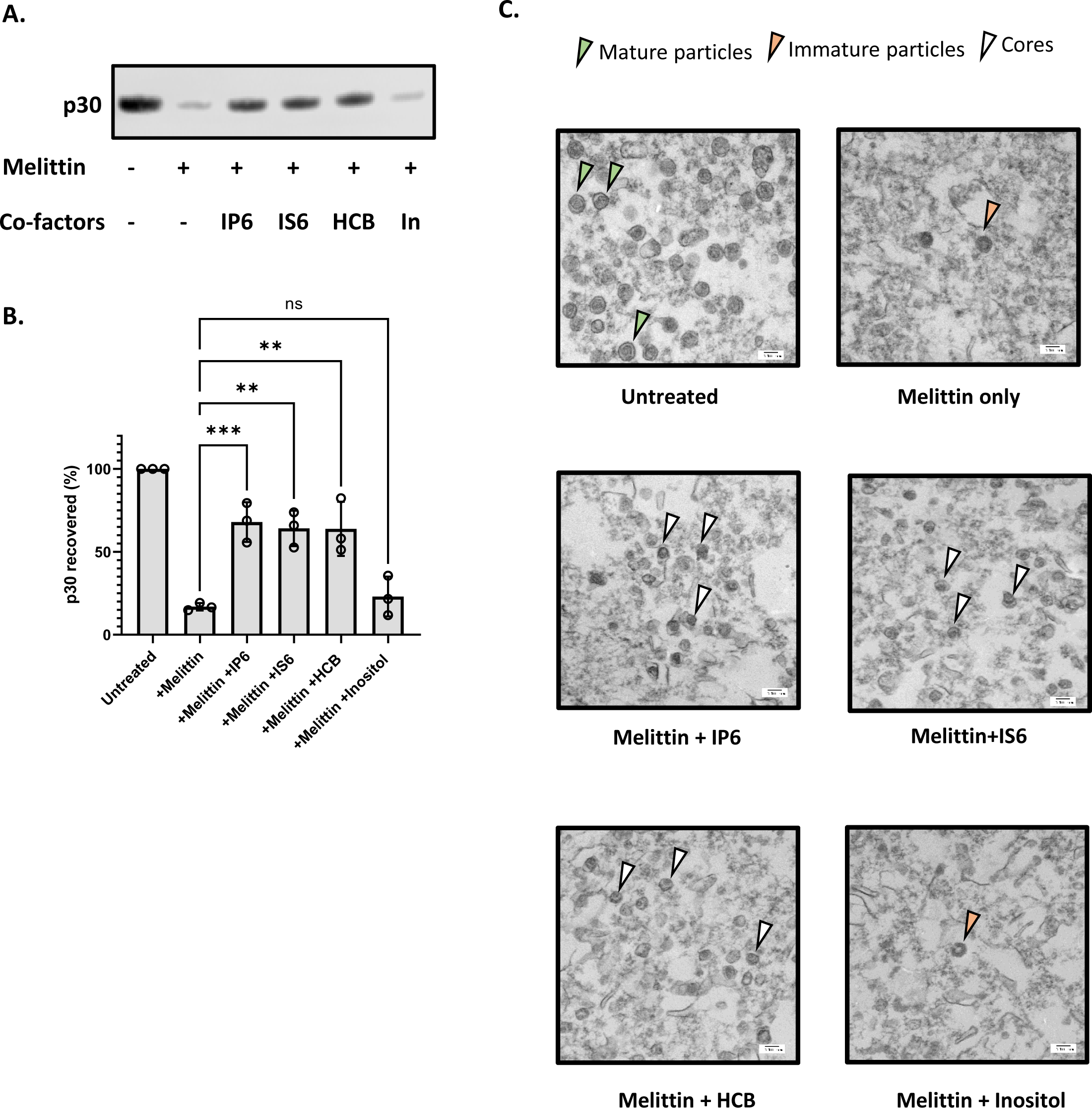
Cyclic polyanions stabilize MLV cores. A) Representative immunoblot probed for MLV capsid protein (p30) in viral pellets in the presence and absence of co-factors (indicated below the blot). The concentration of melittin is 12.5μg/ml and the small molecules are used at a concentration of 80 μM. B) The bar graph shows the quantification of the percentage p30^CA^ recovery after the addition of co-factors. The graph represents the mean ± SD from three independent experiments. Statistical significance is analyzed using one-way ANOVA. P values are indicated by *, ***P<0.001, **P<0.01, ns: not significant. C) Electron micrographs of MLV before and after treatment with co-factors (indicated below each micrograph). The green arrowhead points to mature MLV particles, the orange arrowhead points to immature particles, and the white arrowhead points to intact MLV cores.

### IP6 stabilizes MLV cores

In HIV-1, the stability of the core in permeabilized or lysed virus preparations has a profound effect upon ERT activity, and IP6 appears to promote ERT by stabilizing the cores ^11,20^. We therefore tested the ability of IP6 to stabilize MLV cores. MLV particles were incubated for one hour at 37 °C with melittin in the presence or absence of co-factors tested in Figure 1. The reactions were then centrifuged through a sucrose cushion and the amount of p30^CA^ in the pellet was analyzed by immunoblotting. The results are shown in Figure 2A and the recovery of p30^CA^ in the pellet quantitated in Figure 2B. It is evident that the three active additives, i.e., IP6, IS6, and HCB, all strongly protected the MLV cores from disruption by melittin, as the majority of the CA protein remained pelletable in their presence, but not in inositol (In).

To further characterize the viral components in the ERT reactions, we also examined the pellets by transmission electron microscopy. As shown in Figure 2C, mature viral cores were visible in the pellets obtained after treatment of the particles with melittin and the polyanions IP6, IS6, or HCB. In contrast, the pellets produced in the absence of melittin contained intact mature virions (as expected), while those isolated in melittin alone or with the inactive additive inositol contained few if any mature particles or cores; immature particles were occasionally seen in these samples. Taken together with the immunoblotting results in Figure 2A and 2B, the data indicate that the active additives stabilize the cores within mature MLV particles, in close analogy with prior results on HIV-1^11,20^.

We also tested a wide range of IP6 concentrations in the pelleting assay. Remarkably, the protection against disruption of the cores appeared to be a monotonic function of the IP6 concentration: at 2.4 mM the recovery of p30^CA^ in the pellet was virtually complete (Figure S2A and S2B). This is in striking contrast with the ERT results: as noted above (Figure 1B), IP6 concentrations of 40-80 μM were optimal for ERT, and those above 160 μM were strongly inhibitory. It thus appears that efficient ERT depends upon an intermediate level of capsid stability. Again, these results are quite analogous to previously published findings on HIV-1^11^.

We also tested the ability of rNTPs to stabilize the cores in the pelleting assay. As shown in Figures S2C and S2D, each rNTP was able to stabilize the cores. This is also notable for its contrast with the ERT results, in which (Figure 2) rATP and rCTP were active, while rGTP and rUTP were not. The contrast implies that the enhancement of ERT by rNTPs is not a result of stabilization of the cores.

### MLV packages IP5 and IP6

In light of these striking effects of exogenous IP6 and related molecules on ERT activity, it was of interest to determine whether MLV, like HIV-1 ^3,4^, packages IP6. We prepared MLV particles and assayed them for IP5 and IP6 as described in Star Methods. Control preparations, prepared in parallel with the MLV, included HIV-1 and “mock” virus preparations, which were generated from supernatants of cultures transfected with an empty expression vector. As shown in Figure 3A, we found that IP6 was present in the MLV preparations. As the IP6 levels were significantly higher than that in the “mock” sample, the data strongly indicate that IP6 is an authentic component of MLV particles, rather than a background in the assay. IP5 was also detected in the MLV preparations (Figure 3C). The ratios of IP6 and IP5 to CA protein tended to be somewhat lower in MLV than in HIV-1 (Figure 3B and 3D).

**Figure 3:**
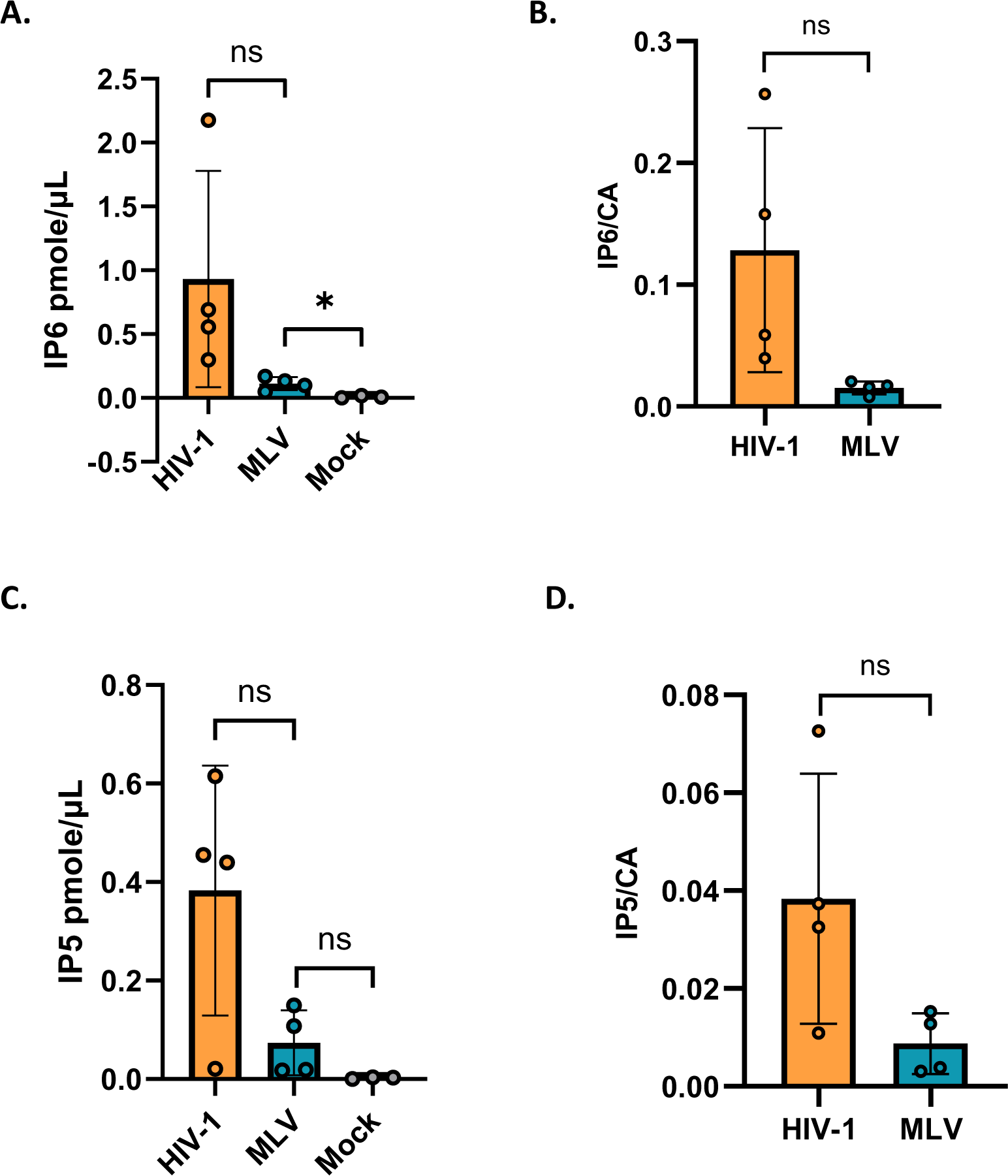
MLV packages IP5/6: The bar graph shows the absolute amount in pmole/µl of A) IP6 and B) IP5 packaged in HIV-1 and MLV. Mock represents supernatant from cells transfected with empty vector control. The amounts of C) IP6 and D) IP5 in HIV-1 and MLV are normalized to the respective capsid protein levels in each virus. The graphs represent the mean ± SD of four independent experiments. Statistical significance is analyzed using Student’s t-test. P values are indicated by *, **P<0.01, *P<0.05, ns: not significant.

### Possible role of R3 residue of MLV capsid in interacting with IP6

In immature HIV-1 particles, IP6 is localized in the centers of Gag hexamers, with a ring of lysines (CA residue 158) above and another ring of lysines (CA residue 227) below the IP6, while in mature particles IP6 is coordinated by a ring of arginines (CA residue 18) in a CA hexamer ^3,4^. It was of interest to try to determine the location of the IP6 within MLV particles. A careful examination of mature MLV using cryo-EM tomography ^23^ previously noted “a faint additional density” within the hexameric N-terminal domain of the CA region of mature particles, surrounded by six arginine side-chains (one from each monomer in the hexamer) of CA residue R3.

It seemed possible that this unidentified density could be IP6. We note that the R3 residue appears to be absolutely conserved in gammaretroviruses (Figure S3A); this conservation suggests that R3 is essential for optimal viral replication and is consistent with a role in packaging of the important cofactors IP5/6. To further test the significance of R3 in MLV, we analyzed the properties of MLV mutants in which it was replaced with lysine or alanine. We found (Figure 4A and 4B) that mature R3K and R3A virions contain IP5 and IP6 levels indistinguishable from those of WT virions; thus, R3 is not necessary for packaging of IP5/6 into virions. On the other hand, we found (Figure 4C) that these mutants lack any detectable infectivity although cells transfected with R3 mutant MLV constructs produced comparable levels of viruses to WT MLV (Figure 4D). In addition, we did not observe any defect in proteolytic processing of Pr65 Gag in the cell lysates of cells transfected with either of the mutants (Figure 4D). We also examined the mutant particles by TEM. We found, just as in WT MLV, two types of virus morphology: one resembling immature MLV, with a ring of density under the virus membrane, and the other with density in the center of the particle, as in mature MLV (Figure S3B). Thus, the mutations did not cause any discernible change in the overall morphology of the particles.

**Figure 4:**
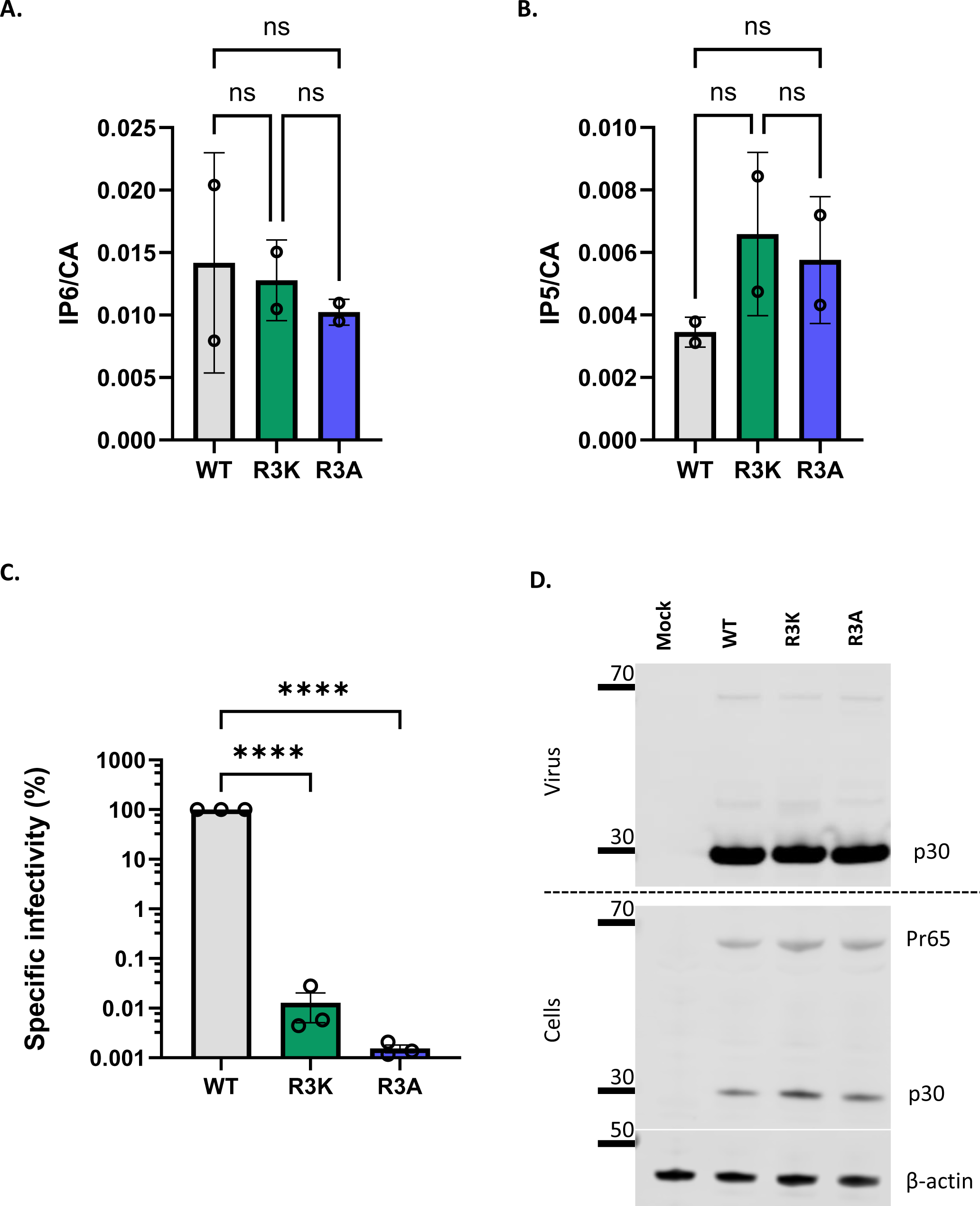
MLV R3 capsid mutants package IP6/5 but are non-infectious. The bar graphs depict the packaging of A) IP6 and B) IP5 within WT, R3K, and R3A MLV viruses. The amount of IP6 and IP5 is normalized to the amount of capsid protein in each of the virus preparations. The graphs represent the mean ± SD from two independent experiments. C) A representative immunoblot against p30^CA^ in the supernatant and cell lysates of cells transfected with either WT MLV or R3 capsid mutants of MLV (R3A and R3K). β-actin is used as a loading control. D) WT, R3A, and R3K MLV plasmids were co-transfected with pBabe-Luc into HEK 293T cells. Viruses produced by these transfected cells were used to infect HT1080mCAT cells, and lysates of these cells were assayed for luciferase activity. Specific infectivity was calculated by normalizing the luciferase values to the MLV p30^CA^ levels (RLU/p30) indicative of the amount of virus in culture supernatants. The graphs represent the mean ± SD of three independent experiments. Statistical significance is analyzed using one-way ANOVA. P values are indicated by *, ****P<0.0001, ns: not significant.

As the mutants were comparable to the WT in their ability to release viruses, we wanted to test if the mutant viral cores are also dependent on IP6 for DNA synthesis *in vitro* using our ERT assay. Interestingly we observed that the R3 mutant virions are apparently incapable of ERT, even in the presence of IP6 (Figure 5A). We also tested the ability of IP6 to stabilize cores from R3 mutant virions; as shown in Figure 5B and 5C, they also differ from WT cores (see Figure 2) in that they are not stabilized by IP6. These results imply that R3 is required for core stabilization by IP6 and are consistent with the hypothesis that this interaction with IP6 is essential for ERT.

**Figure 5:**
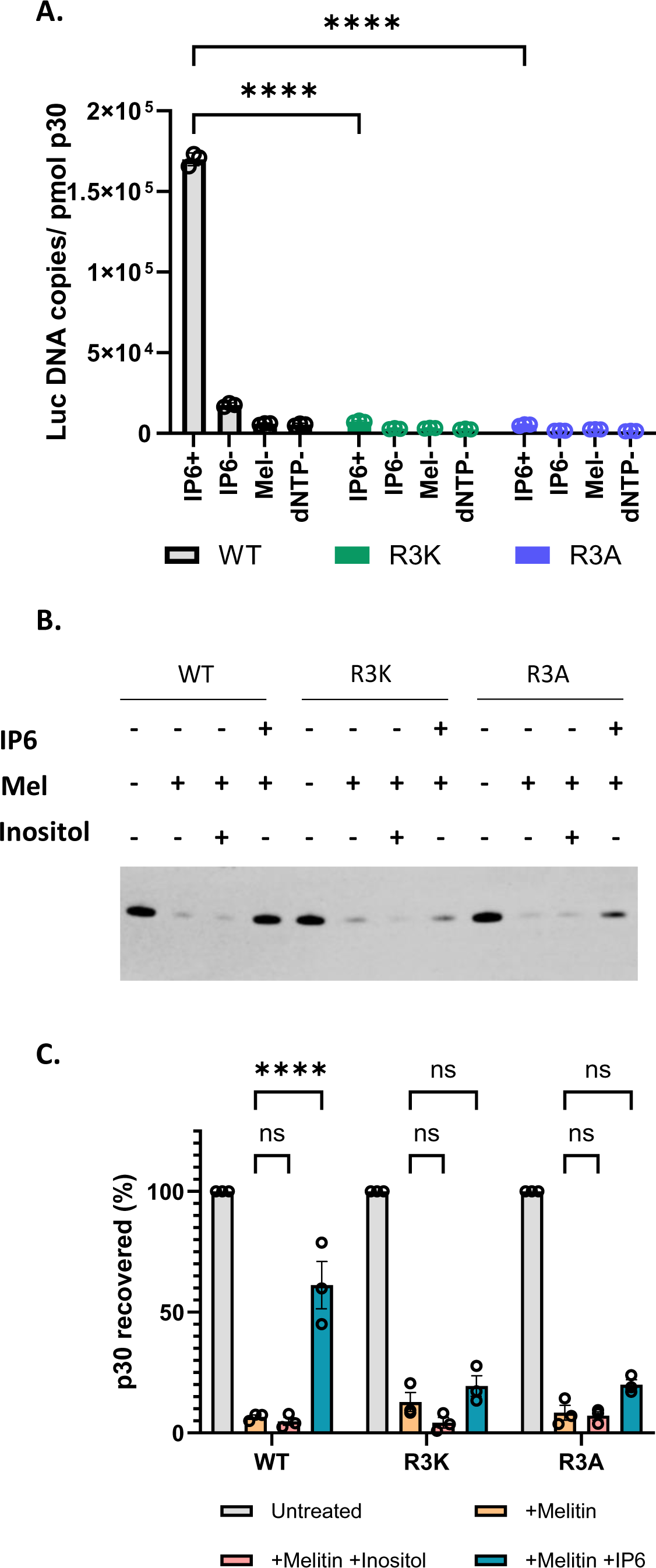
**MLV R3 capsid mutants do not respond to IP6 *in vitro*** A) The bar graph indicates the formation of ERT products in WT MLV and R3 capsid mutants of MLV with or without IP6 as indicated in the x-axis of the graph. The y-axis represents absolute copies of luciferase DNA reverse transcribed from the MLV-derived luciferase vector RNA packaged in MLV normalized to the amount of virus in each sample. B) Shown here is a representative immunoblot that was probed for MLV p30^CA^ in viral pellets derived from WT, R3K, and R3A MLV, under conditions with or without co-factors, as indicated above the blot. The concentration of melittin used was 12.5μg/ml, and co-factors (IP6 and Inositol) were used at 80 μM. C) The bar graph shows the percentage p30^CA^ recovery in pellets after lysis of WT, R3A, and R3K virions in the presence of the indicated co-factors.The graphs represent the mean ± SD from three independent experiments. Statistical significance is analyzed using two-way ANOVA. P values are indicated by *, **** P<0.0001, ns: not significant.

### IP6 functions in MLV replication

While the data presented above demonstrate the effects of IP6 in *in vitro* assays of MLV function, it was obviously of interest to determine the role, if any, of IP6 in viral replication in mammalian cells. We explored this question by testing viral replication in cells that were deficient for IP6 and for IP5. These experiments utilized 293T cells in which the inositol polyphosphate multikinase (IPMK), which synthesizes IP5, or the inositol pentakisphosphate 2-kinase (IPPK), which synthesizes IP6, had been knocked out by CRISPR/Cas9 technology ^24^. It has been shown by two independent studies that upon knockout (KO) of IPMK, levels of both IP5 and IP6 are significantly reduced. However, for the IPPK-KO, IP6 levels were significantly reduced, but IP5 levels were seen to be elevated ^24,25^. We also confirmed these results using direct biochemical assays for IP5 and IP6 and observed that both IPMK- and IPPK-KO cells had lower levels of IP6 than the WT control cells. IPMK-KO cells had a pronounced reduction in IP5 levels, while the IPPK-KO cells only had a modest reduction (Figure S4)

Cultures of the KO and control cells were transfected with plasmids encoding MLV Gag and Gag-Pol; xenotropic MLV Env; and the MLV-based luciferase vector (pBabeLuc). 48 hrs after transfection, the cells were lysed and supernatants were collected for virus assays. Western blot analysis of the cell lysates and supernatants (Figure 6A and Figure S5A, 5B and 5C) showed that expression of viral genes and the rate of virus production were substantially reduced in the KO cells; however, the expressed Gag proteins were assembled into released virions with an efficiency similar to that in the WT control cells (Figure 6B). Significantly, the specific infectivity of the virus released from the KO cells was virtually the same as (or, in the case of the IPMK-KO cells, slightly higher than) that from the control cells (Figure 6C). Thus, normal levels of IP5 and IP6 in virus-producing cells are not necessary for the production of fully infectious MLV.

**Figure 6:**
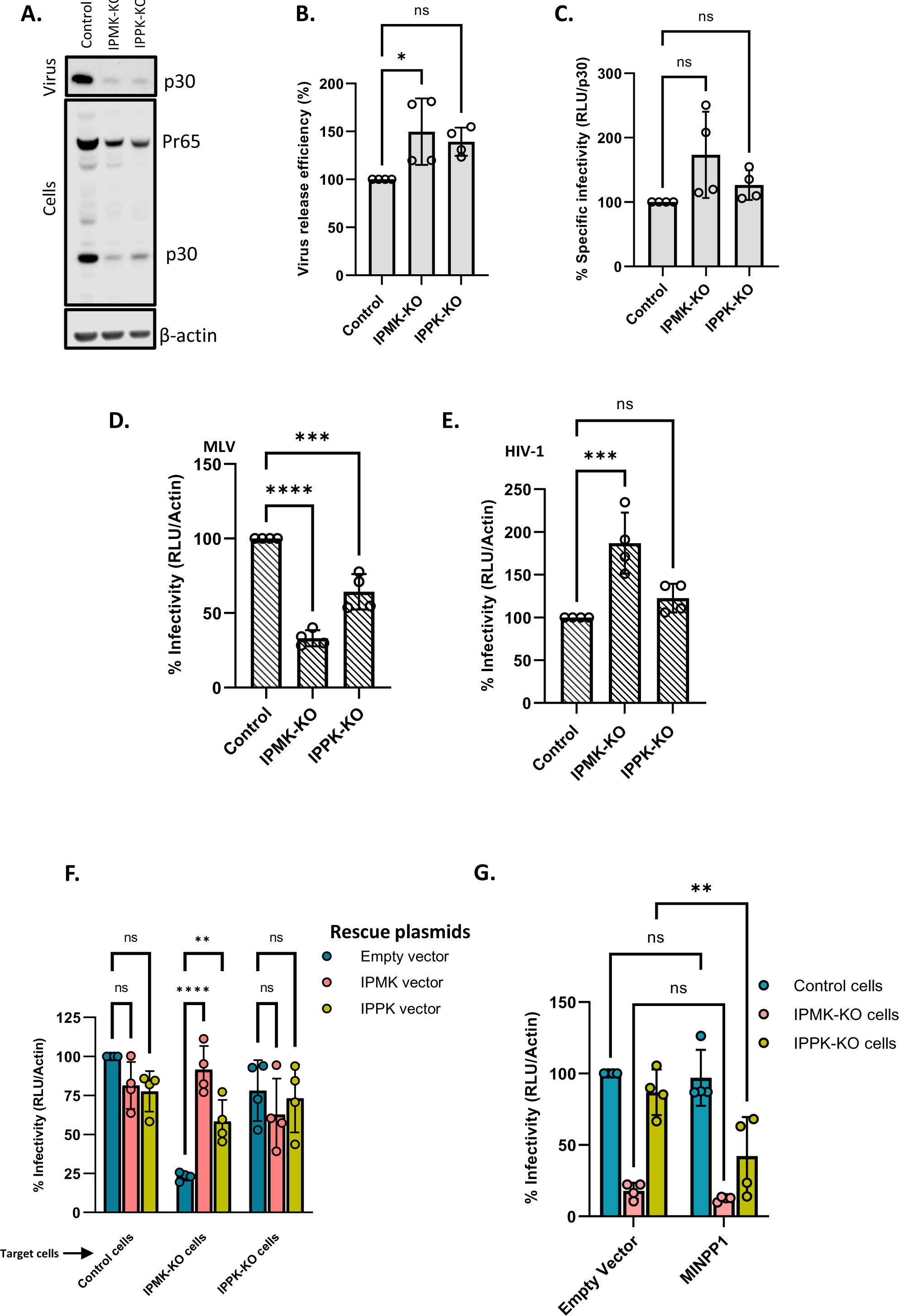
Cellular IP6/IP5 levels contribute to MLV infectivity. A) Control, IPMK-KO, and IPPK-KO cells were transfected with a WT MLV plasmid. The virus was collected from these cultures 48 hours after transfection and both supernatants and cell lysates were analyzed by immunoblotting against p30^CA^. β-actin is used as a loading control. B) Virus release efficiency in the transfected cells. C) **C**ontrol cells, IPMK-KO cells, and IPPK-KO cells were transfected with viral plasmids. Supernatants from these transfected cells were then used to infect control cells and these cells were then assayed for luciferase activity. Specific infectivity was calculated by normalizing the luciferase values to the MLV p30^CA^ levels (RLU/ p30^CA^) . D) Control cells were transfected with Env-defective MLV plasmid (MLV Gag-Pol), Xenotropic MLV Env plasmid, and pBabe-Luc plasmid. Viruses produced by these transfected cells were then used to infect control, IPMK-KO, and IPPK-KO cells and these cells were then lysed and assayed for luciferase activity. Infectivity was calculated by normalizing the luciferase values to the actin levels (RLU/Actin) in the target cells to correct for differences in cell density between control and KO cells. E) Control cells were transfected with HIV-1 pNL4.3ΔEnv (which carries Luciferase in place of the *env* gene) and with the MLV Xenotropic envelope plasmid. Virus from these transfected cells was then used to infect control, IPMK-KO, and IPPK-KO cells and these cells were then lysed and assayed for luciferase activity. Infectivity (RLU/Actin) was measured in control and KO cell lines. The graphs represent the mean ± SD of two independent experiments with two technical replicates in each experiment (n=4). Statistical significance is analyzed using one-way ANOVA. P values are indicated by *, **** P<0.0001, ***P<0.001, *P<0.05, ns: not significant. F) Control and KO cells were transfected with Env-defective MLV plasmid (MLV Gag-Pol), Xenotropic MLV Env plasmid, and pBabe-Luc plasmid with or without IPMK or IPPK expression plasmids. Viruses produced by these transfected cells were then used to infect control cells and these cells were lysed and assayed for luciferase activity. The bar graph shows the rescue of MLV infectivity in KO cells when the missing kinases in the KO cells were added in trans. G) Control cells were transfected with Env-defective MLV plasmid (MLV Gag-Pol), Xenotropic MLV Env plasmid, and pBabe-Luc plasmid. Control and KO cells were then transfected with the MINPP1 expression plasmid or a control plasmid and infected with the virus from the other transfected culture. The cells were lysed and assayed for luciferase activity. The bar graph shows MLV infectivity in control and KO cells transfected with 600 ng of either the empty vector pcDNA3.1 or the MINPP1 expression plasmid. The graphs represent the mean ± SD of two independent experiments with two technical replicates in each experiment (n=4). Statistical significance is analyzed using two-way ANOVA. P values are indicated by *, **** P<0.0001, ** P<0.005, *P<0.05, ns: not significant.

We also tested the possibility that the reduction in IP5 and IP6 levels might render the KO cells less infectable than WT control cells. Parallel cultures of the KO and control cells were infected with virus preparations containing both MLV (with a xenotropic MLV Env) and the MLV-based luciferase vector, and infection was assessed by measuring luciferase activity in the infected cultures. The luciferase activity in the cell lysates was normalized to β-actin levels in the lysates to correct for differences in cell density. We found a highly reproducible reduction in infections in the KO cells (Figure 6D), with a larger decrease in IPMK-KO than in IPPK-KO cells.

It has been reported previously ^5,25^ that IP5/6 deficiency in the target cells does not significantly affect their infectability by HIV-1. Our results with MLV (Figure 6D) are in direct contrast with these findings with HIV-1. To confirm this difference between the viruses, we also checked the infectability of the KO cells with an HIV-1-based luciferase vector. The same MLV Env was used in this pseudotype as in the MLV experiments shown in Figure 6D. As shown in Figure 6E, the KO of IPMK and of IPPK did not reduce the efficiency with which the cells could be infected with HIV-1, in agreement with prior studies^5,25^.

The data presented above indicate that KO cells produce normal, fully infectious MLV, while the efficiency with which they can be infected by MLV is somewhat reduced. As an additional test of this conclusion, we tested the ability of MLV produced from KO cells to infect the KO cells. As shown in Figure S5D and S5E, this virus exhibits the same profile as the virus produced in WT cells (see Fig. 6D): a modest but highly reproducible reduction in infections in the KO cells, particularly the IPMK-KO cells. This confirms that the reduced IP5/6 levels in the virus-producing cells have no significant effect upon the dependence of the virus upon IP5/6 levels in the target cells.

To further confirm that the reduced infection of the KO cells resulted from the lack of the IPMK and IPPK, we tested whether transient transfection with expression vectors for the missing enzymes could reverse the effect. As shown in Figure 6F, the defect in infectability of the IPMK-KO cells was almost completely reversed by either of the kinases, while there was little effect in the IPPK-KO cells, whose infectability is much closer to that of the control cells than that of the IPMK-KO cells.

We also tested the ability of another enzyme, multiple inositol polyphosphate phosphatase (MINPP1), to reduce the infectability of target cells. This enzyme converts IP6 to IP5 and IP5 to IP4, reducing cellular levels of both IP6 and IP5 ^26,27^. Using the same methodology as in Figure 6F, we transfected the control and KO cells with an expression plasmid for MINPP1 and challenged them with the MLV preparations containing the luciferase vector. As shown in Figure 6G, we found that this treatment reduced infection of the IPPK-KO cells, while we could not detect an effect in the control cells or the IPMK-KO cells. As part of this experiment, we also checked the expression of MINPP1 in the transfected cells by immunoblotting. Interestingly, we found (Figure S6) that the enzyme was at a substantially higher level in the IPMK-KO cells than in the IPPK-KO cells or control cells. This unexpected observation raises the possibility that MINPP1 expression in 293T cells is controlled by IP5 or IP6 levels, but we have not explored this further. In any case, the fact that MINPP1 reduced the infectability of IPPK-KO cells further supports the conclusion that infection by MLV depends upon the maintenance of proper IP5 and/or IP6 levels in target cells.

### Reduced infectability of KO cells is reflected in reduced viral DNA synthesis

We also tested whether the rate or amount of viral DNA synthesis is reduced upon infection of the KO cells. The KO and control cells were infected with MLV preparations (containing both intact MLV and the luciferase vector) and were lysed at 3, 6, and 10 hours after infection. The lysates were assayed for early, intermediate, and late DNA products, as well as luciferase DNA, by qPCR. They were also assayed for the cellular gene CCR5, which was used to normalize the lysates for differences in cell numbers. As shown in Figure 7, in all cases reverse transcription was impaired in the KO cells. The defect was most severe in late DNA synthesis and was greater in the IPMK-KO cells than in the IPPK-KO cells, mirroring the relative infectability of the cells (Figure 6D).

**Figure 7:**
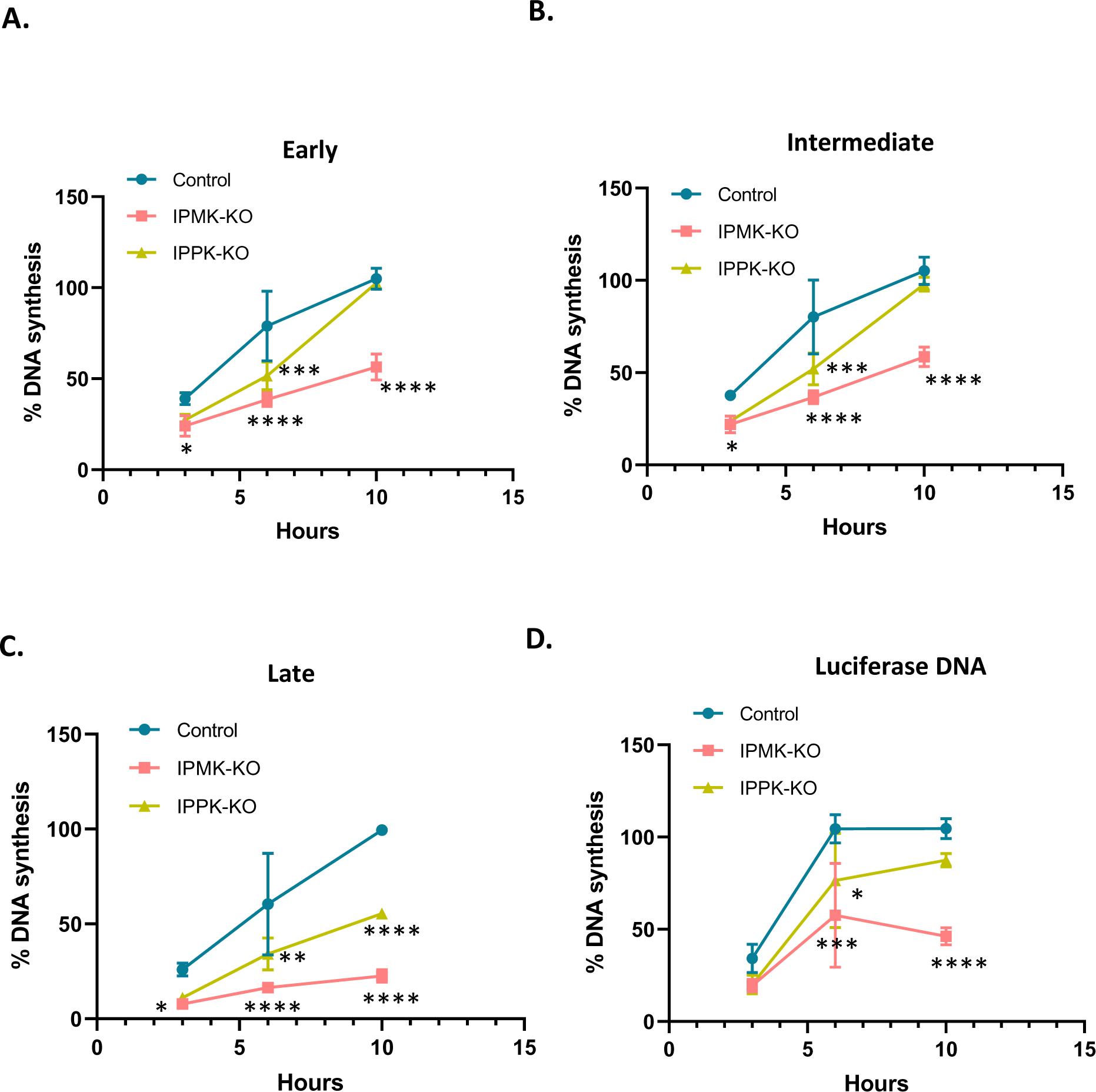
IP6/5 is required for reverse transcription during MLV infection. Percentage MLV DNA synthesis measured in infected cells (control and IP-KO cells) using qPCR with primers for A) Early B) Intermediate and C) late reverse transcription products. Before infection, virus preparations were treated with DNase to remove plasmid DNA, as described in Materials and Methods. DNA was extracted from cell lysates at indicated time points within the first 10 hours after infection. D) As some of the viral particles contained pBabeLuc-derived RNA, DNA synthesis was also measured using luciferase primers as in ERT assays. DNA copies of MLV and luciferase were normalized to host CCR5 copies to account for differences in cell density; in each graph, “100” represents the value obtained at 10 hours after infection in the control cells. The graphs represent the mean ± SD of two independent experiments with two technical replicates in each experiment (n=4). Statistical significance is analyzed using two-way ANOVA. P values are indicated by *, **** P<0.0001, ***P<0.001, ** P<0.005, *P<0.05, ns: not significant.

## DISCUSSION

In the last few years, the small molecule IP6 has been found to play a significant role in HIV-1 replication ^7^. It is present at substantial levels within virions. It both promotes assembly of the immature particle, and enhances the stability of the core within mature virions. This stabilization appears to facilitate DNA synthesis within the core in newly infected cells ^11^.

In the present work, we have investigated the participation of IP6 in the replication of MLV, a member of the gammaretrovirus genus. Our results can be briefly summarized as follows: just as in HIV-1, IP6 is packaged in MLV particles. In assays of ERT (the copying *in vitro* of viral RNA to DNA by the RT in viral lysates), we found that adding IP6 profoundly enhances DNA synthesis. Also in analogy with prior work on HIV-1 ^11,20^, this enhancement reflects the stabilization of the structure of the viral core by exogenously supplied IP6 in particles permeabilized by the pore-forming peptide melittin or, alternatively, lysed by the mild nonionic detergent Triton X-100. Finally, cell-based infection assays reveal that MLV infection depends on the presence of IP6 and IP5 in target cells, while in the case of HIV-1, such dependence is absent.

Our analysis of the contribution of IP6 to MLV replication was made possible by the existence of cell lines from which the two biosynthetic enzymes, IPPK and IPMK, had been (separately) knocked out^24^. Briefly, we found that when IPPK-KO and IPMK-KO cells were transfected with plasmids encoding infectious MLV, they produced virus with unimpaired specific infectivity, albeit at a lower level than the control cultures. This is analogous to previously published findings on HIV-1 ^24,25^.

On the other hand, infection of IPMK-KO or IPPK-KO target cells with normal MLV was somewhat less efficient than infection of controls; this is clearly contrary to results with HIV-1 ^5,25^, which we confirmed as part of our study. It is interesting to note that the defect in infectability was greater in the IPMK-KO cells than the IPPK-KO cells, that the low infectability of the IPMK-KO cells could be reversed by transient transfection of plasmids encoding either IPMK or IPPK into the cells, and also that the infectability of the IPPK-KO cells was further reduced by transient transfection of a plasmid encoding MINPP1, a phosphatase that reduces the phosphorylation of IP6 and related compounds ^26,27^. (The experiments with MINPP1 also strongly suggested that steady-state levels of this enzyme in 293T cells may be negatively regulated in response to IP5, but we have not explored this observation further.) This target-cell effect is the one fundamental difference we have found between MLV and HIV-1 with respect to IP6 function. We consider its significance below.

Several observations in this study should also be noted. In addition to IP5 and IP6, the small cyclic polyanions IS6 and HCB also promote ERT in MLV. While HCB and IP5 are somewhat less active on a molar basis than IP6, IS6 is nearly equivalent to IP6 in this assay. This is rather unexpected as IS6 carries significantly less negative charge than IP6. In any case, these charges are apparently essential for promotion of ERT, as inositol, the uncharged parent of IP6 and IS6, appears to be completely devoid of activity (Figure 1).

We also found that rNTPs are necessary for the ERT reaction. One might imagine that they serve the same function in the reaction as IP6, as they are also small polyanions, but this does not seem to be the case: our experiments gave no indication that there was redundancy in these requirements. In fact, further analysis showed that the “rNTP” requirement can be fulfilled by rATP or rCTP, but not, as far as we could determine, by rGTP or rUTP (Fig. S1). We do not know how these specific nucleoside triphosphates contribute to ERT, but it is particularly surprising that there is base-specificity in the reaction and that of the two active rNTPs, one has a purine and the other a pyrimidine base. The contribution of rNTPs to ERT also cannot be explained by an ability to stabilize the core, as all four rNTPs possessed this activity (Fig. S2), but, as noted here, only rATP and rCTP supported ERT. In contrast, in HIV-1 it has been found that either rATP or rGTP can protect disulfide-stabilized CA hexamers against thermal denaturation ^4^; rATP also enhances ERT in HIV-1 ^4^.

Our experiments also showed that IP6 imparts some degree of stability to cores isolated from MLV virions and that this stability seems necessary for the synthesis of viral DNA *in vitro*. On the other hand, supraoptimal levels of IP6 apparently produce hyperstable cores which cannot synthesize DNA efficiently, in further resemblance to HIV-1 ^11^. It seems likely that just as with HIV-1 ^11,20^, addition of IP6 will be very helpful in the purification of MLV cores from mature virions.

We also addressed the question of the location of IP6 within mature MLV cores. Qu et al. ^23^ had reported that a ring of six arginines at residue 3 of MLV CA appears to coordinate an unidentified density; it seemed possible that this density is IP6. We tested the significance of this residue by replacing it with lysine or with alanine. These mutants possess no detectable infectivity (Fig. 4) and are evidently incapable of ERT, even in the presence of IP6 (Fig. 5). IP6 also fails to stabilize their cores in melittin-disrupted particles (Fig. 5). All of these findings are consistent with the hypothesis that IP6 is coordinated by R3 in mature MLV cores, although other explanations cannot be excluded. We also found that these mutant virions still contain IP6 (Fig. 4); thus, R3 is not necessary for IP6 packaging.

Why should wild-type levels of IP5 and/or IP6 in the target cells be required for successful infection by MLV, when this is not the case for HIV-1? One key difference in the replication of the two viruses is that unlike HIV-1, MLV cannot infect non-dividing cells ^13,28^. Recent studies have shown that in HIV-1, the core of the mature infecting particle remains partly or entirely intact while it is in the cytoplasm and copying its RNA into DNA ^16–18,29,30^. The core is capable of penetrating the nucleus by interacting with cellular nuclear-import machinery, and only dissociates and releases the viral DNA (complexed with integrase [IN]) within the nucleus. In MLV, in contrast, the DNA cannot penetrate the interphase nucleus. Rather, the core at least partially disassembles within the cytoplasm, and a complex of DNA and IN, together with the Gag fragment p12 as well as CA, gains access to nuclear DNA by binding to mitotic chromosomes in dividing cells ^12–15^.

We propose that because MLV cores disassemble in the cytoplasm during infection, they are dependent upon the cellular IP6 for maintaining the structural stability required for efficient reverse transcription. Cores of HIV-1 particles, however, bring IP6 with them and never relinquish it during their journey to the nucleus. Perhaps HIV-1 cores have a higher affinity for IP6 than MLV cores. In any case, this would explain why the reduction of IP6 levels in target cells does not impair HIV-1 infection. The nature of the “trigger” leading to the disassembly of MLV cores, but not HIV-1 cores, in the cytoplasm of newly infected cells remains to be determined.

## ACKNOWLEDGEMENTS

We would like to acknowledge Eric. O. Freed and Leo James for providing the IP6/5 KO cell lines. The NCI’s electron microscopy core facility, particularly Ferri Soheilian, contributed TEM images. We thank Dr. John York for sharing plasmids and advice regarding enzymatic assays of IP5 and IP6. This research was supported in part by the NIH Intramural Research Program, National Cancer Institute, Center for Cancer Research, and the NIH Intramural AIDS Targeted Antiviral Program. Additional funding was provided through NIH extramural research grant R21 AI150384.

## AUTHOR CONTRIBUTIONS

Conceptualization, B.B., K.K.L., S.K.D and A.R.; Methodology, B.B., K.K.L., H.B., S.K.D., D.H., G.A.S., C.A., and A.R.; Investigation, B.B., K.K.L., H.B., S.K.D., D.H., C.A., and A.R; Writing – Original Draft, B.B and A.R,; Writing – Review & Editing, A.R.; Funding Acquisition, A.R. and C.A.; Resources, A.R. and C.A.; Supervision, S.K.D., C.A., and A.R.

## DECLARATION of INTERESTS

The authors declare no competing interest.

## STAR METHODS

### Cells and viruses

#### MLV production for ERT assays

All cell lines used in this study were maintained in Dulbecco’s modified Eagle’s medium (DMEM) supplemented with 10% fetal bovine serum (Hyclone), 100 U/ml of penicillin, and 100 µg/ml of streptomycin, and were grown at 37°C with 5% CO_2_. MLV for ERT assays was produced from HEK293T cells by co-transfecting 5μg of pNCS plasmid (a full-length MLV genome, a kind gift from Stephen Goff, ^31^), and 5μg pBabe-Luc plasmid (a pBabe-puro-derived luciferase vector ^32^) in a 10cm dish using TransIT 293 (Mirus Bio). (Transfecting these plasmids together will give rise to virions with WT MLV proteins; some will contain the full-length, replication-competent MLV genome and some will contain the pBabeLuc genome.) Medium was changed ∼18h post-transfection. Virus-containing supernatant was collected 48h post-transfection, pooled from several plates, and cleared by filtration through a 0.45µ filter. The filtered virus-containing supernatant was pelleted through a 20% (w/v) sucrose cushion, prepared in 1X HS buffer (10mM Hepes, pH 7.4, 140mM NaCl) in an SW28 rotor (Beckman) at 25000 rpm for 2 hours. The viral pellet in each tube was resuspended in 150µl of 1X HS buffer and stored at -80°C for use in ERT assays (ERT virus). The amount of p30^CA^ in the pellet was quantitated using p30 ELISA (Cell Biolabs) according to the manufacturer’s protocol.

#### MLV production for infectivity assays

For infectivity assays, MLV was produced from three different cell lines 1) control cells, which are HEK293T cells; 2) IPMK-KO HEK293T cells, where the IPMK gene had been knocked out ^24^; and 3) IPPK-KO HEK293T cells, where the IPPK gene had been knocked out ^24^. All three cell lines were a gift from Leo James. MLV was produced in control cells and the two KO cells in parallel, by co-transfecting 5µg of Env-defective MLV plasmid (which also encodes Glyco-Gag ^33^) (MLV Gag-Pol), 1 µg of Xenotropic MLV Env plasmid (a kind gift from Henrich Gottlinger ^34^), and 5µg of pBabe-Luc in a 10 cm dish. The R3 MLV capsid mutant viruses (R3K and R3A) were produced from HEK293T cell line by co-transfecting 5μg of pNCS plasmid containing the intended mutation and 5μg pBabeLuc plasmid in a 10cm dish using TransIT 293 (Mirus Bio). Medium was changed ∼18 hours post-transfection. Virus-containing supernatant was collected 48h post-transfection and cleared by filtration through a 0.22µ syringe filter. Filtered supernatants were stored at -80°C.

### Reverse transcription assays

To assess ERT, 100 µl of ERT virus aliquots were first digested with DNase I (TURBO DNA-free™ Kit, ThermoFisher Scientific) to remove transfected DNA contamination. DNase I digestion was carried out in a total volume of 200µl at 37°C for 30 mins followed by treatment with DNase I inactivation resin supplied with the kit according to the manufacturer’s instructions. The treated virus was then used for ERT assay. Each ERT reaction was set up in a total volume of 20µl containing 25mM Tris pH 7.4, 75mM NaCl, 60µM dCTP, 46µM dGTP, 80µM dTTP, 52µM dATP, 0.18mM rCTP, 1.75mM rGTP, 0.7mM rUTP, 6.7mM rATP, 3.3 mM MgCl_2_, 40 µM IP6, 6.25 µg/ml melittin (Sigma-Aldrich) and 8 µl of DNase-treated ERT-virus unless otherwise specified. The concentrations of Tris, rNTPs, and MgCl_2_ were similar to those used in Christensen et al. ^11^, while the dNTP concentrations were 10-fold higher than those in Christensen et al. ^11^. The reaction was incubated at 37°C for 10 hours for the synthesis of reverse-transcribed DNA products. DNA products were column-purified using Nucleospin Gel and PCR clean-up kit (Machery-Nagel) and eluted in 25ul of elution buffer (Machery-Nagel). The reverse-transcribed luciferase DNA was quantitated using SYBR Green 1-based qPCR with FastStart Essential DNA Green Master (Roche Life Sciences) and primers luc PB F and luc PB R (Key Resources Table) that targeted the luciferase coding sequence. For measuring ERT kinetics, the ERT reaction was halted at time points specified in the figures: the reaction mixture was immediately applied to the column for purification of the products and quantification using qPCR. Primers used for the quantification were either specific to (-) strand strong stop (MSSF4, MSSR2, ^35^) for early products; sequences synthesized after the first-strand transfer (MFST-F, MFST-R) for intermediate products; or those made after second-strand transfer for late products (M2ST-F, M2ST-R). The target sequences for these primers are present in both pNCS (full-length MLV genome) and pBabe-Luc to enable the detection of the total amount of DNA products formed from both MLV RNA and pBabe-Luc RNA. Quantification of copies from qPCR was done using a standard curve generated with every assay using serial dilutions of pBabeLuc plasmid. Ct values of test samples were within the range of the standard curve. To amplify the full luciferase coding sequence (CDS), primers Luc34F and Luc1525R were used ^36^. Sequences of all primers are given in the Key Resources Table.

ERT activity for R3 capsid mutants (R3K and R3A) was assessed as above. The ERT products (luciferase DNA) from WT, R3K and R3A measured using qPCR were normalized to the amount of virus, which was quantified using MLV p30 ELISA (Cell Biolabs).

To assess whether IP6 affects the enzymatic activity of MLV RT, we measured RT activity in lysed virions using the SG-PERT assay ^37^ as follows. A 50 µl aliquot of ERT virus was treated with 50 µl 2X-PERT lysis buffer containing 0.25% Triton X-100, 50mM KCl, 100mM Tris-Cl pH7.4, and 40% glycerol. These virus lysates were then added to 40 ng of MS2 RNA (Sigma Aldrich), primers (MS2-F and MS2-R), 2X ERT buffer without rNTPs, and different concentrations of IP6, and incubated at 37°C for 50 mins for reverse transcription. Standards were made by serial dilution of recombinant MMLV RT (ThermoFisher Scientific) and incubating with an external template and primers parallel to test samples. Reverse-transcribed products were column-purified and quantitated by SYBR green-based qPCR using primers MS2-F and MS2-R.

### Core stability assay

To assess the stability of MLV cores in the presence of small molecules, ERT-virus preparations were used. The standard reaction contained, where indicated, 12.5 µg/ml of melittin; 80 µM of IP6, IS6, HCB, or inositol; and/or rNTPs at 6.7 mM each. The reaction was incubated at 37°C for 1 hour and then centrifuged through a 20% sucrose cushion in an SW 55 Ti rotor (Beckman) at 28000 rpm for 1.5 hours. Pelleted material was resuspended in 100µl 1X NuPAGE LDS sample buffer (Invitrogen) containing NuPAGE sample reducing agent (Invitrogen) and 1X HALT protease inhibitor and analyzed for p30 by immunoblotting.

To measure RT activity in the pellets, the pelleted material was resuspended in 100µl of 1X PERT lysis buffer, diluted 10-fold with a buffer comprised of 25 mM KCl + 50 mM Tris HCl pH 7.5, and assayed using the SG-PERT procedure described above.

### Electron microscopy

The effects of small molecules upon the stability of viral cores were also analyzed using electron microscopy. After samples were incubated in melittin and small molecules at 37°C for 1 hour as described above, they were fixed by adding 2% glutaraldehyde solution. The fixed samples were then centrifuged in a beem capsule and the pellets were stained, sectioned, and visualized by transmission electron microscopy; details of methods are available on request.

### Enzymatic assays of inositol phosphates

IP5 and IP6 were quantified by enzymatic conversion to radiolabeled IP6 and IP7 in reactions catalyzed by purified recombinant Ipk1 and VIP2, respectively, in the presence of ATP-[γ-32P]. The enzymes were expressed in E. coli and purified as described ^38^. Products were separated by thin-layer chromatography and detected and quantified by phosphorimager analysis. IP5 reactions were performed in 10-µL volumes containing 50 mM Tris (pH 8.0), 10 mM MgCl_2_, 62 ng/µL GST-atIpk1, with 10 µCi ATP-[γ-32P] (6,000 Ci/mmol, 10 mCi/mL, PerkinElmer Life Sciences). IP6 assay reactions were performed in 10-µL volumes containing 50 mM Bis-Tris (pH 6.0), 10 mM MgCl_2_, 62 ng/µL GST-hsVIP2, and 10 µCi of ATP-[γ-32P]. Reactions of standards contained 25 fmol to 0.4 fmol of IP5 (Cayman Chemical) or 0.02 pmol to 0.62 pmol IP6 (TCI). Prior to addition to the reactions, virus samples were lysed by addition of Triton X-100 to 0.1% (vol/vol) and heated at 95°C for 10 minutes to dissociate bound inositol phosphates from proteins.

Ipk1-and VIP2-catalyzed reactions were incubated at 37°C for 60 min. To stop the reactions, 0.5 µL of 20 mg/mL proteinase K was added to each sample and incubated at 56°C for 10 min. The samples were then incubated at 70°C for 10 min to heat-inactivate the proteinase K. 2.5 µl of each sample was applied to polyethyleneimine-cellulose thin layer chromatography (TLC) plates (Millipore Sigma 105579) that were pre-dried in an oven at 60°C for at least an hour. The spots were then allowed to air-dry at room temperature for 10 min and were developed in a TLC tank equilibrated with fresh elution buffer consisting of 1.09 M KH_2_PO4, 0.72 M K_2_HPO4, and 2.26 M HCl or 2.5 M HCl for IP5 and IP6 reactions, respectively. Plates were dried at 60°C for 20 min. Plates were exposed to a phosphor storage screen that was subsequently scanned on a FLA7000IP Typhoon phosphorimager (GE Healthcare). Images were quantified using Image Studio Lite (LI-COR Biosciences) and metabolites were quantified by interpolation of nonlinear curve fitting by second-order polynomial using GraphPad Prism 9 from values that fell within the useful range of the standards.

### Immunoblotting

For specific infectivity assays, MLV p30^CA^ was quantified directly from the virus-containing supernatant using rabbit polyclonal anti-p30^CA^ antiserum. The supernatant was diluted in NuPAGE LDS sample buffer (Invitrogen) containing NuPAGE sample reducing agent (Invitrogen) and 1X HALT protease inhibitor (ThermoFisher Scientific). The virus-producing cells were also prepared using the same NuPAGE LDS sample buffer cocktail, sonicated for complete lysis, and probed as above. For pelleting-based assays, pelletable material was also assayed using the anti-p30^CA^. Samples to be assayed were boiled at 90°C for 5 mins, loaded onto NuPage 4-12% Bis-Tris PAGE, and electrophoresed until the dye reached the bottom of the gel. Separated proteins were then transferred to Immobilon-FL transfer membranes (Millipore).

Membranes were blocked using the Intercept (TBS) blocking buffer from LI-COR. After blocking, membranes were probed overnight at 4°C with the rabbit anti-p30^CA^ antiserum diluted in blocking buffer. Membranes were washed with TBS (20mM Tris, pH 7.0, 500mM NaCl) buffer before incubation at room temperature for 1 hour with IRdye800 donkey anti-rabbit secondary antibody (LI-COR Biosciences). Membranes were imaged on Odyssey system (LI-COR Biosciences) followed by quantification of the bands using Image Studio Lite (LI-COR Biosciences). Cell lysates were also probed for β-actin using mouse anti-β-actin (Abcepta). For detecting MINPP1 in MINPP1-overexpressing cells, cell lysates were probed with anti-MINPP1 antiserum (Fabgenix). In assays measuring infectivity in target cells the amount of β-actin in cell lysates was also quantified. For quantification of band intensities for p30 and β-actin, a standard curve was prepared using dilutions of a test sample run on the same gel. All sample band intensities fell within the linear range of the standard curve.

### Virus Quantitation in IP6 Packaging Assays

Quantification of CA proteins in preparations of HIV-1 particles was performed by immunoblotting of viral lysates in parallel with purified recombinant HIV-1 CA (purified according to Ganser et al. ^39^) and MLV CA-NC protein (purified as described ^40^). The concentrations of the purified reference proteins were determined spectrophotometrically using extinction coefficients that were calculated according to the method of von Hippel ^41^. The concentrations were confirmed by protein assay using the bicinchoninic acid method (BCA Protein Assay kit, ThermoFisher). Values obtained by the two methods agreed to within 5%. Samples were subjected to electrophoresis on 4-20% gradient polyacrylamide gels containing SDS (Genscript). Proteins were transferred electrophoretically to nitrocellulose membrane, and the blots were probed with HIV-1 CA-specific monoclonal antibody 183-H12-5C and goat anti-AKR p30^CA^ polyclonal antiserum obtained from the NCI/BCB Repository located at ViroMed Biosafety Laboratories. Following probing with IR dye-conjugated secondary antibodies, the bands were detected with a LI-COR Odyssey imager and quantified using the instrument software. Values for the viral samples were interpolated from standard curves generated from the signals for the recombinant proteins. The preparations were analyzed in three independent experiments, and the mean values were employed in calculating IP6:CA stoichiometry.

### Infectivity assays

In all infections, cells were pre-treated with medium containing DEAE-dextran (20 μg/ml) for 30 minutes before exposure to the virus-containing supernatant, which had been diluted 10-fold in complete medium. This inoculum was left on the cells for 48 hrs and the cells were then lysed for luciferase assays, as in the manufacturer’s instructions (Promega)

Control 293T cells were infected with viruses produced from control, IPMK-KO, and IPPK-KO cells to measure specific infectivity. Cells were lysed 48h post-infection and assayed for luciferase activity. Specific infectivity was calculated by normalizing luciferase activity to the amount of the virus. The amount of virus was quantified by immunoblotting for p30 in the filtered virus-containing supernatant. Virus release efficiency was calculated using the formula described in Mallery. et. al. for HIV-1 ^24^, i.e.,

(𝑣𝑖𝑟𝑢𝑠 𝑝30)/(𝑣𝑖𝑟𝑢𝑠 𝑝30 + 𝑐𝑒𝑙𝑙 𝑝30 + 𝑐𝑒𝑙𝑙 𝑃𝑟65)

For assessing the effect of IP6/5 depletion in target cells upon MLV replication, infectivity assays were performed in which an equal volume of virus-containing supernatant from the control cells was used to infect control, IPMK-KO cells, and IPPK-KO cells as target cells. Infectivity was assayed by measuring luciferase activity in the cell lysates 48 h post-infection and normalizing the values to β-actin levels in the lysates, as determined by immunoblotting. For rescue experiments, control cells, IPMK-KO cells, and IPPK-KO cells were first transfected with 500 ng of IPMK or IPPK expression plasmid (a kind gift of Leo James). After 24 hours the transfected cells were infected with MLV. Cell lysates were assayed as above. Similarly, the effect of MINPP1 expression upon infection was assessed by transfecting control or KO cells with 600 ng of MINPP1 plasmid; the cells were infected 24 h after transfection and harvested for luciferase and actin assays 48 h after transfection.

Infectivity assays for the R3 mutants were performed in the HT1080-mCAT cell line ^33^ using the same procedure described above.

### MLV DNA synthesis in infected cells

The reverse transcription products in MLV-infected cells were assayed as follows. Cells were initially infected (following pre-treatment with DEAE-dextran as described above) with virus that had been DNase digested using DNase I, RNase Free (Invitrogen) to remove plasmid DNA contamination. An aliquot of the DNase I-treated supernatant was heat-inactivated at 68°C for 20 mins. These pre-treated virus-containing supernatants were added to control cells, IPMK-KO cells, and IPPK-KO cells. Cells were lysed 3 hours, 6 hours, and 10 hours post-infection and the DNA was extracted using a QIAamp DNA mini kit (Qiagen). Extracted DNA was quantified using SYBR green-based (FastStart Essential DNA Green Master, Roche Life Sciences) qPCR for early, intermediate and late products using the same primers used for assaying kinetics in ERT assays, or for luciferase DNA using the luciferase primers used in the ERT assay. DNA copies were also measured in cells infected with the heat-inactivated virus after DNaseI treatment to assess the effectiveness of the DNaseI treatment. Luciferase DNA copies in cells infected with heated virus were approximately 100-fold lower than in cells infected with unheated viruses. To account for differences in the recovery of DNA from cells, MLV DNA copy numbers measured with each of the primers were normalized to the copy numbers of the cellular gene CCR5 quantified using primers CCR5-For and CCR5-Rev ^32,42^.

### Statistical analysis

The statistical significance of all the data was either analyzed by Student’s t-test, one-way, or two-way analysis of variance (ANOVA) using GraphPad Prism 10. The figure legends state the statistical test used in each figure.

## KEY RESOURCES TABLE

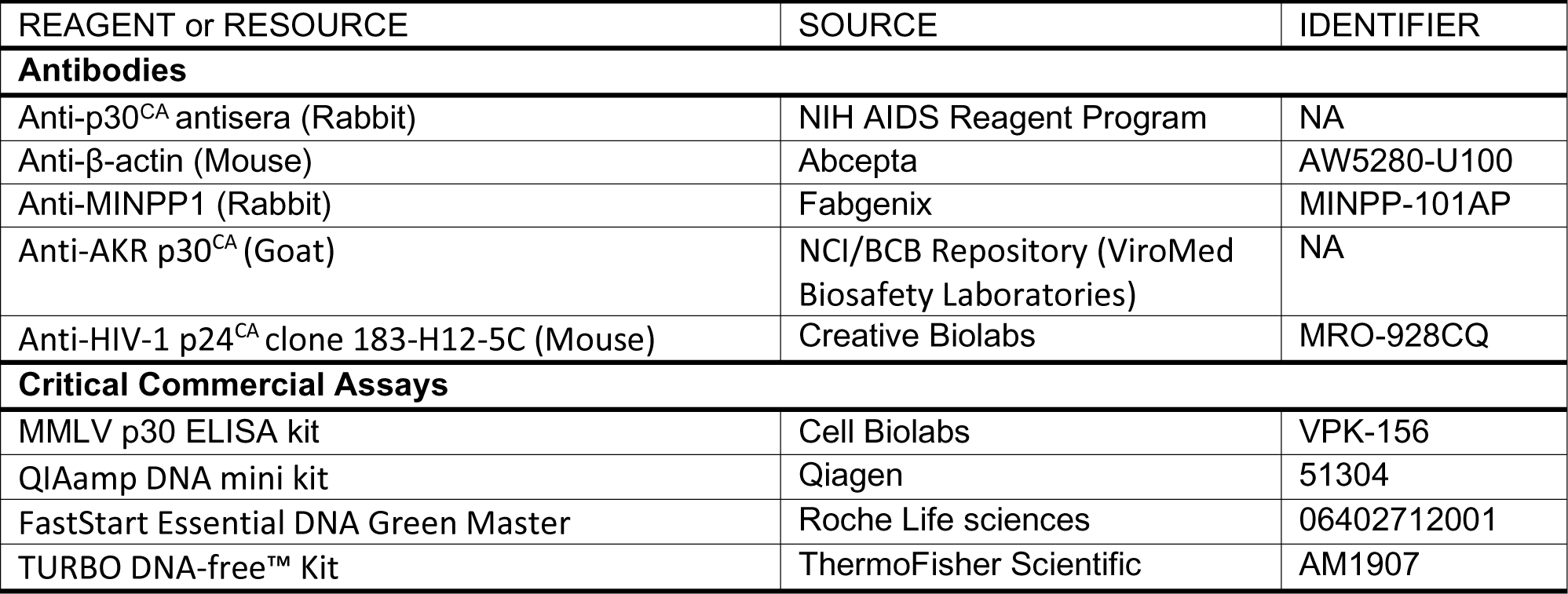
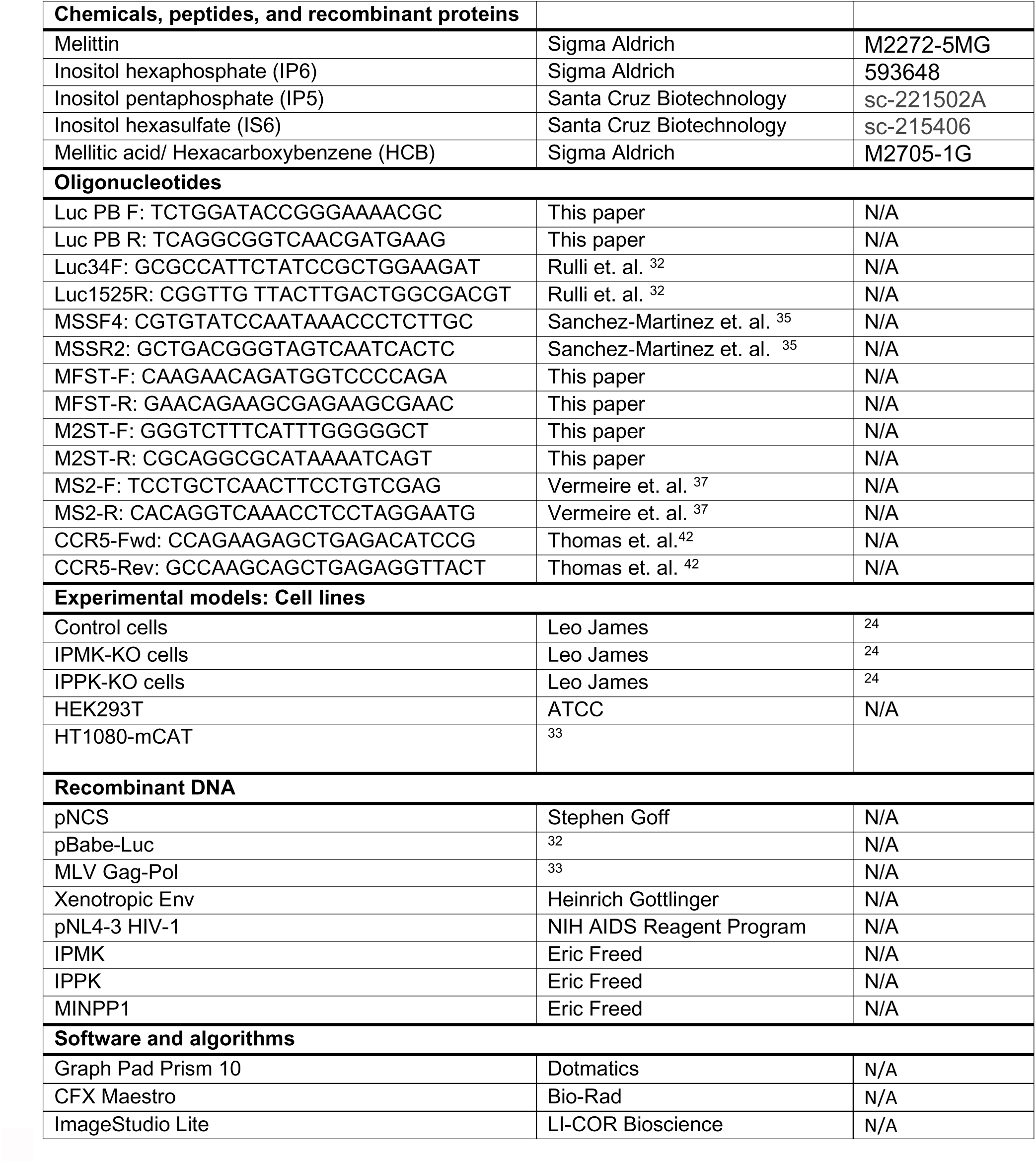

## SUPPLEMENTAL INFORMATION TITLES AND LEGENDS

Figure S1: **Optimization of ERT reaction**: A) The bar graph indicates ERT product formation in the presence of different concentrations of melittin (x-axis). B) ERT product formation was also observed when Triton-X was used instead of melittin. IP6 concentration in these assays was 40µM. C) Early, intermediate, and late ERT product formation measured during different time intervals (x-axis). D) The bar graph shows the formation of ERT products in different reaction conditions indicated below the graph. Concertation of the rNTP mixture is the same as in the ERT buffer (described in materials methods) unless mentioned otherwise in the X-axis. The label “6.7mM” above three bars refers to the concentration of each rNTP. E) IP6 titration with and without rNTPs in the ERT reaction buffer. The graphs represent the mean ± SD of three replicates in the qPCR measurement of a single experiment. F) The bar graph shows MLV RT activity measured using SG-PERT assay. Viruses were lysed in PERT lysis buffer containing 0.125% of triton-X for near-complete lysis. Lysed samples were incubated with an external template (MS2 RNA) and primers. Reverse transcribed products were measured using qPCR. The graphs represent the mean ± SD of three replicates in the qPCR measurement of a single experiment. Except for Figure S1E, all other figures are representative of a single experiment selected from two independent experiments.

Figure S2: **Pelleting-based stability assays:** A) Immunoblot showing recovery of MLV capsid protein with increasing concentration of IP6. Inositol is used as a negative control at a concentration of 80µM. B) Quantification of the p30^CA^ recovery from the immunoblot. C) Immunoblot showing MLV capsid protein (p30^CA^) recovery after the addition of rNTPs (6.7mM each). IP6 was used as a positive control and inositol was used as a negative control, both at 80µM. D) Quantification of percent p30^CA^ recovery from immunoblots of three independent experiments. The graphs represent the mean ± SD of three independent experiments. Statistical significance is analyzed using one-way ANOVA. P values are indicated by *, ***P<0.001, ** P<0.01, *P<0.05, ns: not significant.

Figure S3: **R3 residue in MLV capsid is conserved and MLV-WT and R3 capsid mutant particles have similar morphology:** A) Sequence alignment of various gammaretrovirus capsid regions shows complete conservation of the R3 residue, which is highlighted in red. The following gammaretroviruses are included in the alignment: Moloney murine leukemia virus (MMLV, Acc No. P03355), Friend virus (FV, Acc No. P26808.2), Amphotropic murine leukemia virus (A-MLV, Acc No. AAO61195.1), Xenotropic murine leukemia virus (XMRV, Acc No. A1Z651.1), Moloney murine leukemia virus neuropathogenic variant (MLVMN, Acc No. Q8UN02.2), endogenous ecotropic MLV in AKR mice (AKV, Acc No. P03356.3), Feline leukemia virus (FeLV, Acc No. NP_047255.1), Koala retrovirus (KoRV, Acc No. Q9TTC1.2), Porcine endogenous retrovirus (PERV, Acc No. CAB65339.1), Gibbon ape leukemia virus (GaLV, Acc No. P21414.2), and RD114 retrovirus (a cat endogenous virus) from a human tumor cell line RD, Acc No. BAM17305.1. Conservation annotation is as follows: an asterisk (*) denotes positions with a single, fully conserved residue, a colon (:) indicates conservation among groups of strongly similar properties, and a period (.) signifies conservation among groups of weakly similar properties. B) Transmission electron microscopy (TEM) analysis of WT and R3 capsid mutants of MLV. HEK293T cells were transfected with full-length MLV-WT or R3A and R3K mutants for virus production. Two days later they were processed for thin-section TEM. The green arrowhead indicates MLV particles displaying electron density underlying the virion membrane, while the red arrowhead points to MLV particles exhibiting electron density in the center of the virion. The scale bar is 200nm.

Figure S4: **IP6/5 analysis in IPMK-KO and IPPK-KO cells**: A) TiO2 PAGE showing levels of IP6 in control, IPMK-KO, and IPPK-KO cells. B) The bar graph shows the amount of IP6 and IP5 quantitated from TLC in control, IPMK-KO, and IPPK-KO cells.

Figure S5: **IP6 is required for MLV replication:** Quantitation of MLV p30^CA^ in the supernatant (A); MLV p30^CA^ (B), and Pr65 (C) in the cell lysates of control cells and KO cells shown in the representative immunoblot of Figure 5A. Infectivity (RLU/Actin) in control cells vs KO cells with viruses produced from D) IPMK-KO and E) IPPK-KO. Infectivity was calculated after normalizing the luciferase values to the actin levels in the target cells to account for differences in cell density between the control and KO cells. The graphs represent the mean ± SD of two independent experiments with two technical replicates in each experiment (n=4). Statistical significance is analyzed using one-way ANOVA. P values are indicated by *, **** P<0.0001, ***P<0.001.

Figure S6: **MINPP1 expression in control and IP-KO cells:** Immunoblot showing MINPP1 expression in the control and the KO cell lines transfected with 200 ng or 600 ng of MINPP1-expressing plasmid or of pcDNA3.1.

## Notes

### Competing Interest Statement

The authors have declared no competing interest.

