## Supplementary Information for "Essential functions of Inositol hexakisphosphate (IP6) in Murine Leukemia Virus replication"

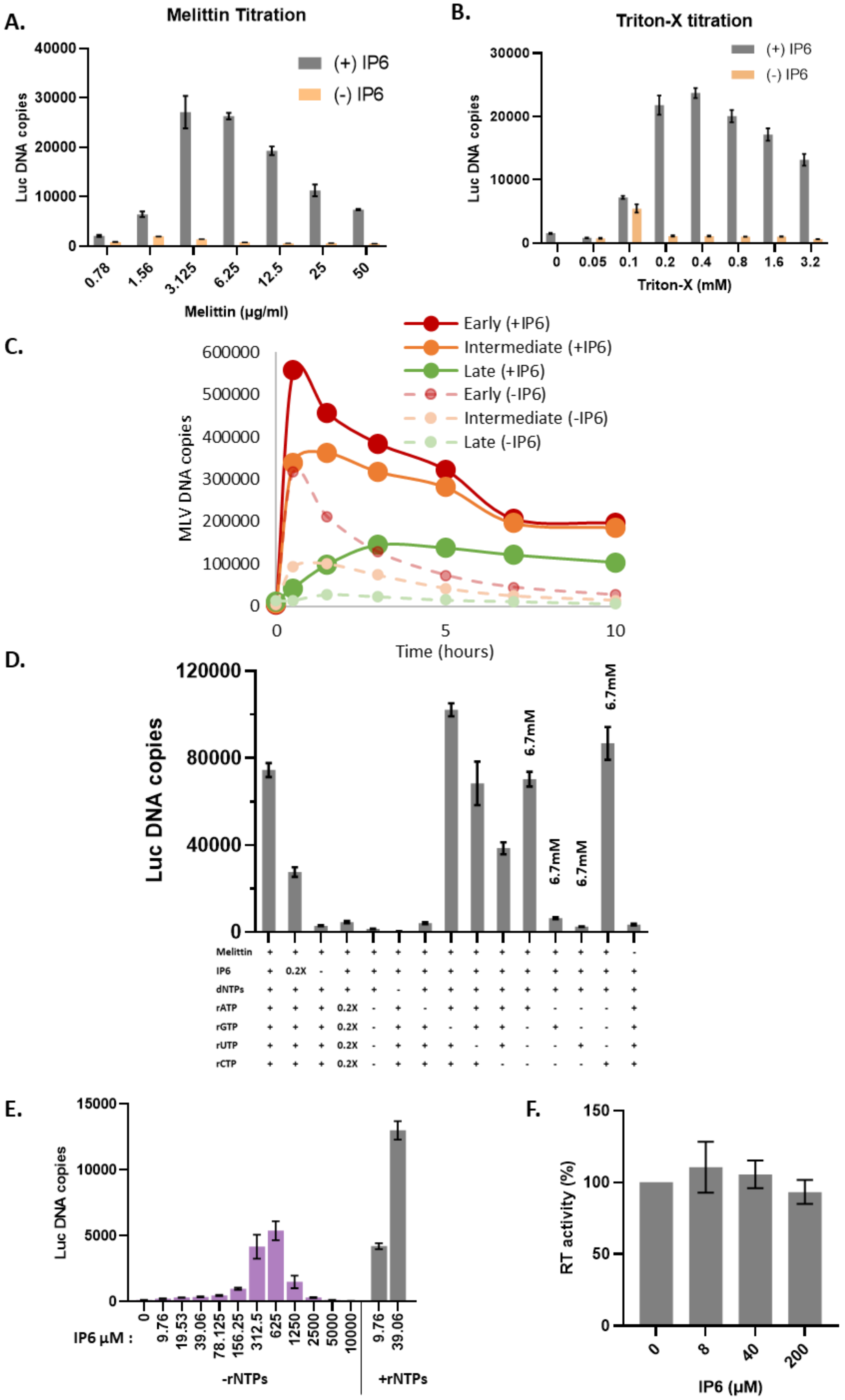
A.
Melittin Titration
(+) IP6
(-) IP6
Luc DNA copies
0
10000
20000
30000
40000
0.78
1.56
3.125
6.25
12.5
25
50
Melittin (μg/ml)
B.
Triton-X titration
(+) IP6
(-) IP6
Luc DNA copies
0
10000
20000
30000
0
0.05
0.1
0.2
0.4
0.8
1.6
3.2
Triton-X (mM)
C.
Early (+IP6)
Intermediate (+IP6)
Late (+IP6)
Early (-IP6)
Intermediate (-IP6)
Late (-IP6)
MLV DNA copies
0
100000
200000
300000
400000
500000
600000
0
5
10
Time (hours)
D.
Luc DNA copies
0
40000
80000
120000
6.7mM
6.7mM
6.7mM
6.7mM
Melittin + + + + + + + + + + + + + + -
IP6 + 0.2X - + + + + + + + + + + + +
dNTPs + + + + + - + + + + + + + + +
rATP + + + 0.2X - + - + + + + - - - +
rGTP + + + 0.2X - + + - + + - + - - +
rUTP + + + 0.2X - + + + - + - - + - +
rCTP + + + 0.2X - + + + + - - - - + +
E.
Luc DNA copies
0
5000
10000
15000
IP6 μM :
0
9.76
19.53
39.06
78.125
156.25
312.5
625
1250
2500
5000
10000
9.76
39.06
-rNTPs
+rNTPs
F.
RT activity (%)
0
50
100
150
0
8
40
200
IP6 (μM)

**Figure S1: Optimization of ERT reaction**: A) The bar graph indicates ERT product formation in the presence of different concentrations of melittin (x-axis). B) ERT product formation was also observed when Triton-X was used instead of melittin. IP6 concentration in these assays was 40µM. C) Early, intermediate, and late ERT product formation measured during different time intervals (x-axis). D) The bar graph shows the formation of ERT products in different reaction conditions indicated below the graph. Concertation of the rNTP mixture is the same as in the ERT buffer (described in materials methods) unless mentioned otherwise in the X-axis. The label "6.7mM" above three bars refers to the concentration of each rNTP. E) IP6 titration with and without rNTPs in the ERT reaction buffer. The graphs represent the mean ± SD of three replicates in the qPCR measurement of a single experiment. F) The bar graph shows MLV RT activity measured using SG-PERT assay. Viruses were lysed in PERT lysis buffer containing 0.125% of triton-X for near-complete lysis. Lysed samples were incubated with an external template (MS2 RNA) and primers. Reverse transcribed products were measured using qPCR. The graphs represent the mean ± SD of three replicates in the qPCR measurement of a single experiment. Except for Figure S1E, all other figures are representative of a single experiment selected from two independent experiments.

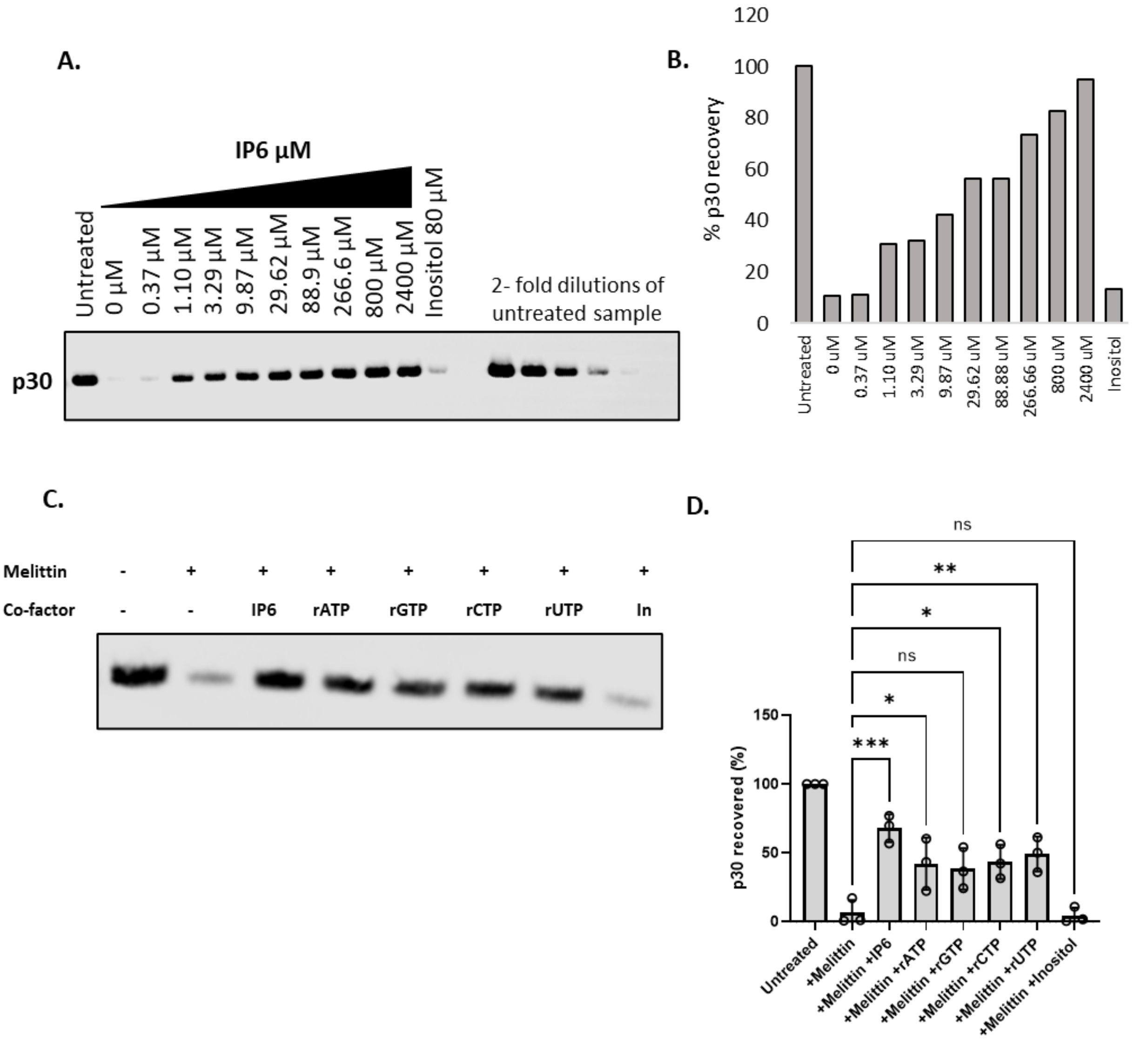


**Figure S2: Pelleting-based stability assays:** A) Immunoblot showing recovery of MLV capsid protein with increasing concentration of IP6. Inositol is used as a negative control at a concentration of 80μM. B) Quantification of the $p30^{CA}$ recovery from the immunoblot. C) Immunoblot showing MLV capsid protein ($p30^{CA}$) recovery after the addition of rNTPs (6.7mM each). IP6 was used as a positive control and inositol was used as a negative control, both at 80μM. D) Quantification of percent $p30^{CA}$ recovery from immunoblots of three independent experiments. The graphs represent the mean ± SD of three independent experiments. Statistical significance is analyzed using one-way ANOVA. P values are indicated by *, ***P<0.001, ** P<0.01, *P<0.05, ns: not significant.

**A.**

```
MMLV         1 PLRAGGN--------GQLQYWPFSSSDLYNWKNNNPSFSEDPGKLTALIESVLITHQPTWDDCQQLLGTLLTGEE  67
FV           1 PLRQGGN--------GQFQYWPFSSSDLYNWKNNNPSFSEDPAKLTALIESVLLTHQPTWDDCQQLLGTLLTGEE  67
A-MLV        1 PLRSGGN--------GQLQYWPFSSSDLYNWKNNNPSFSEDPGKLTALIESVLLTHQPTWDDCQQLLGTLLTGEE  67
XMRV         1 PLRMGGD--------GQLQYWPFSSSDLYNWKNNNPSFSEDPGKLTALIESVLITHQPTWDDCQQLLGTLLTGEE  67
MLVMN        1 PLRTGGN--------GQLQYWPFSSSDLYNWKNNNPSFSEDPGKLTALIESVLITHQPTWDDCQQLLGTLLTGEE  67
AKV          1 PLRLGGN--------GQLQYWPFSSSDLYNWKNNNPSFSEDPGKLTALIESVLTTHQPTWDDCQQLLGTLLTGEE  67
FeLV         1 PLREGPN--------NRPQYWPFSASDLYNWKSHNPPFSQDPVALTNLIESILVTHQPTWDDCQQLLQALLTGEE  67
KoRV         1 PLRAVGPPAEPNGL-VPLQYWPFSSADLYNWKSNHPSFSENPTGLTGLLESLMFSHQPTWDDCQQLLQVLFTTEE  74
PERV         1 PLRTYGPPM-PGGQLQPLQYWPFSSADLYNWKTNHPPFSEDPQRLTGLVESLMFSHQPTWDDCQQLLQTLFTTEE  74
GaLV         1 PLRAIGPPAEPNGL-VPLQYWPFSSADLYNWKSNHPSFSENPAGLTGLLESLMFSHQPTWDDCQQLLQILFTTEE  74
RD114        1 PLRTV-N--------RTVQYWPFSASDLYNWKTHNPSFSQEPQALTSLIESILLTHQPTWDDCQQLLQVLLTTEE  66

Conservation   ***               ******::******.::*.**::*  ** *:**:: :************  *:* **
```

**B.**

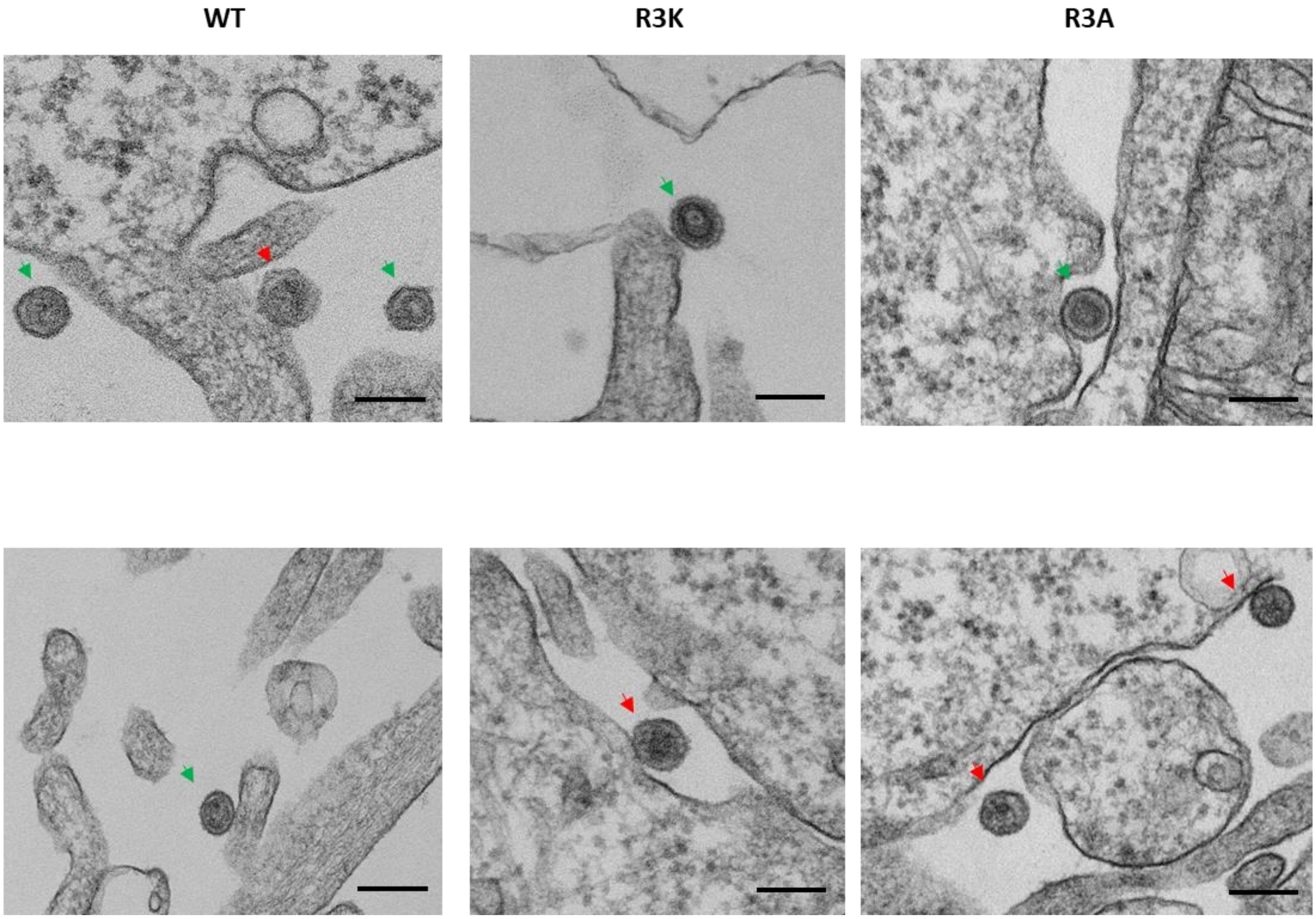

**Figure S3: R3 residue in MLV capsid is conserved and MLV-WT and R3 capsid mutant particles have similar morphology:** A) Sequence alignment of various gammaretrovirus capsid regions shows complete conservation of the R3 residue, which is highlighted in red. The following gammaretroviruses are included in the alignment: Moloney murine leukemia virus (MMLV, Acc No. P03355 ), Friend virus (FV, Acc No. P26808.2 ), Amphotropic murine leukemia virus (A-MLV, Acc No. AAO61195.1), Xenotropic murine leukemia virus (XMRV, Acc No. A1Z651.1 ), Moloney murine leukemia virus neuropathogenic variant (MLVMN, Acc No. Q8UN02.2), endogenous ecotropic MLV in AKR mice (AKV, Acc No. P03356.3), Feline leukemia virus (FeLV, Acc No. NP_047255.1), Koala retrovirus (KoRV, Acc No. Q9TTC1.2), Porcine endogenous retrovirus (PERV, Acc No. CAB65339.1), Gibbon ape leukemia virus (GaLV, Acc No. P21414.2), and RD114 retrovirus (a cat endogenous virus) from a human tumor cell line RD, Acc No. BAM17305.1. Conservation annotation is as follows: an asterisk (*) denotes positions with a single, fully conserved residue, a colon (:) indicates conservation among groups of strongly similar properties, and a period (.) signifies conservation among groups of weakly similar properties. B) Transmission electron microscopy (TEM) analysis of WT and R3 capsid mutants of MLV. HEK293T cells were transfected with full-length MLV-WT or R3A and R3K mutants for virus production. Two days later they were processed for thin-section TEM. The green arrowhead indicates MLV particles displaying electron density underlying the virion membrane, while the red arrowhead points to MLV particles exhibiting electron density in the center of the virion. The scale bar is 200nm.

**A.**

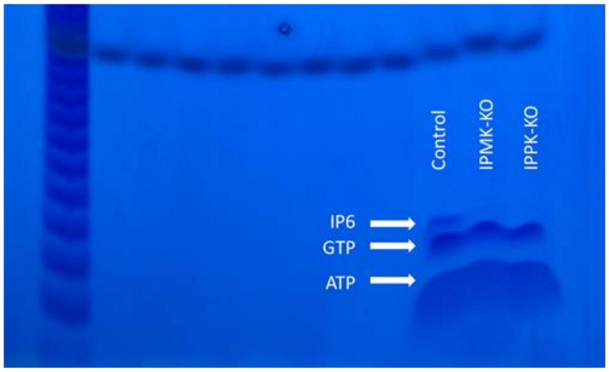


**B.**

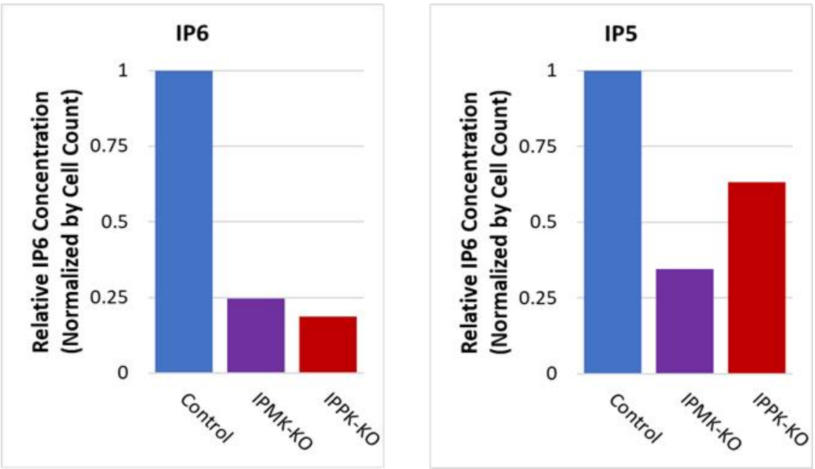

**Figure S4: IP6/5 analysis in IPMK-KO and IPPK-KO cells**: A) TiO2 PAGE showing levels of IP6 in control, IPMK-KO, and IPPK-KO cells. B) The bar graph shows the amount of IP6 and IP5 quantitated from TLC in control, IPMK-KO, and IPPK-KO cells.

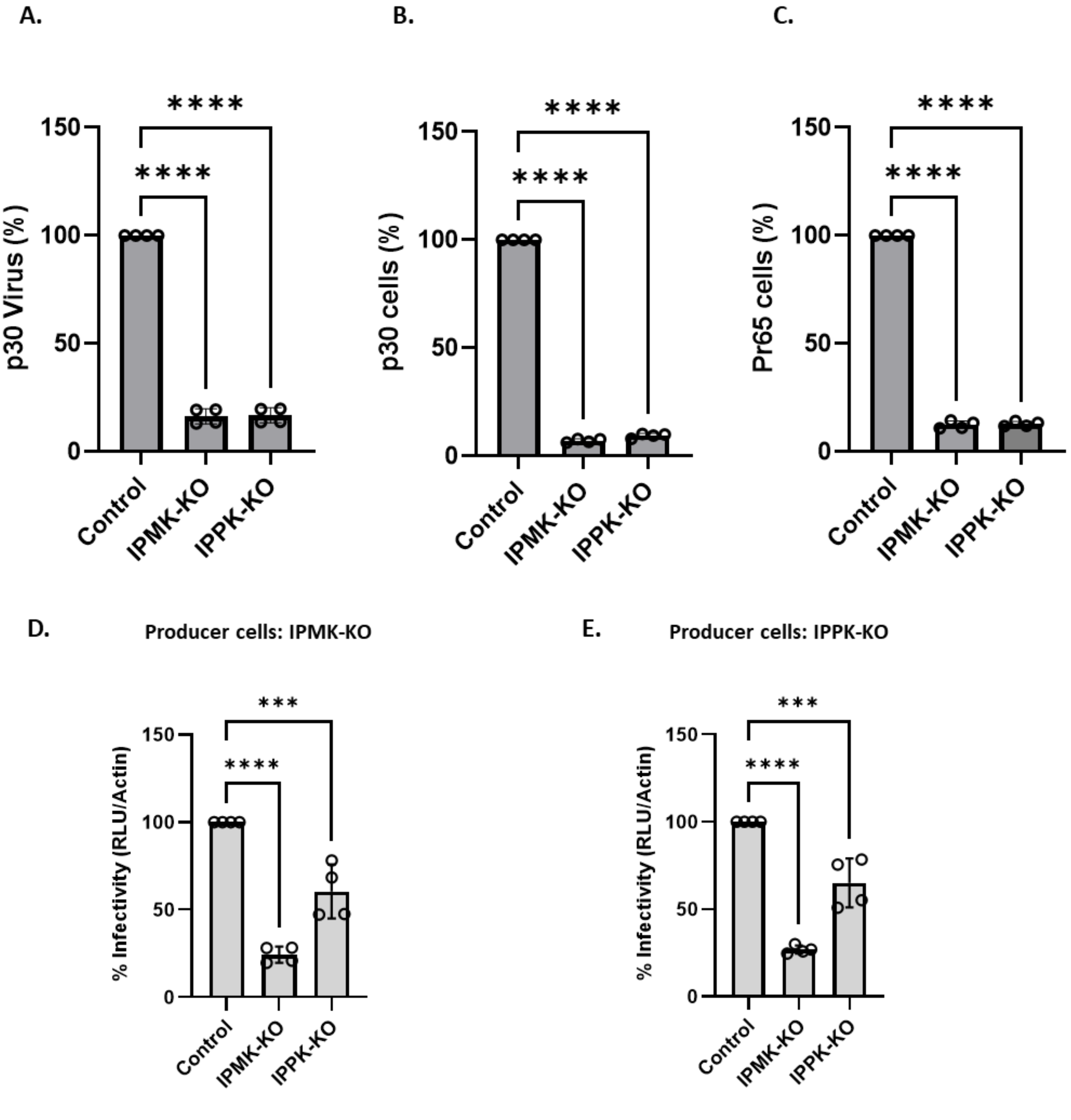


**Figure S5: IP6 is required for MLV replication:** Quantitation of MLV $p30^{CA}$ in the supernatant (A); MLV $p30^{CA}$ (B), and Pr65 (C) in the cell lysates of control cells and KO cells shown in the representative immunoblot of Figure 5A. Infectivity (RLU/Actin) in control cells vs KO cells with viruses produced from D) IPMK-KO and E) IPPK-KO. Infectivity was calculated after normalizing the luciferase values to the actin levels in the target cells to account for differences in cell density between the control and KO cells. The graphs represent the mean ± SD of two independent experiments with two technical replicates in each experiment (n=4). Statistical significance is analyzed using one-way ANOVA. P values are indicated by *, **** P<0.0001, ***P<0.001.

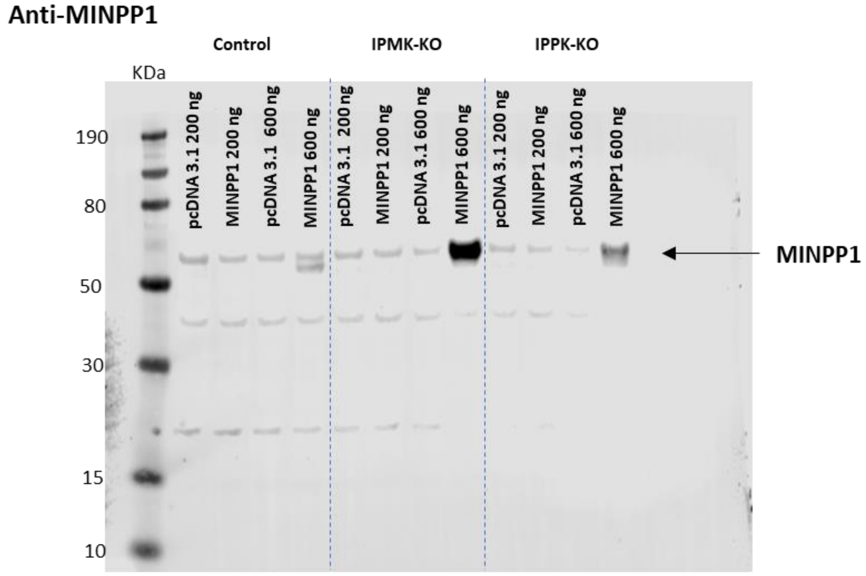


**Figure S6: MINPP1 expression in control and IP-KO cells:** Immunoblot showing MINPP1 expression in the control and the KO cell lines transfected with 200 ng or 600 ng of MINPP1-expressing plasmid or of pcDNA3.1.
